# Evaluating features of scientific conferences: A call for improvements

**DOI:** 10.1101/2020.04.02.022079

**Authors:** Sarvenaz Sarabipour, Aziz Khan, Samantha Seah, Aneth D. Mwakilili, Fiona N. Mumoki, Pablo J. Sáez, Benjamin Schwessinger, Humberto J. Debat, Tomislav Mestrovic

## Abstract

Scientific conferences provide valuable opportunities for researchers across career stages and disciplines to present their latest work, and to network with their peers. These meetings have largely been held in-person with rapid proliferation in the number of meetings and attendees. Yet the format and quality of their organization lag behind what is possible and as a result, the current experience of attending conferences in many disciplines remains unchanged in many respects. We created a database of 270 national and international academic conferences held in-person during 2018-2019 in various disciplines and examined them for their features, costs and impact on the community. We found that many meetings could still be improved significantly in terms of diversity, inclusivity, promoting early career researcher (ECR) networking and career development, venue accessibility, and importantly, reducing the meetings’ carbon footprint. It is important to accelerate and mandate efforts to improve conferences so that researchers in all disciplines, in particular ECRs, consistently benefit from scientific gatherings, for years to come. We discuss a combination of approaches and recommendations to make conferences more modern, effective, equitable and intellectually productive for the research community and environmentally sustainable for our planet.

*“They always say time changes things, but you actually have to change them yourself*.*”*

*— Andy Warhol*

## Introduction

Scientific conferences are important avenues for researchers to share and discuss research findings, to exchange ideas and insights, and to network for collaboration and career development. Organizing inclusive and useful scientific meetings is a significant responsibility and a service to the research community. This requires passion, dedication, considerable time and thoughtful planning and is shared by researchers, scientific societies, and other organizations worldwide. There has been a sharp increase in the size of the academic workforce and in the number of scientific meetings organized (1–3) yet, after over 180 years, most conferences are still held in person and provide limited attendance opportunities for many researchers from diverse socio-economic backgrounds, particularly early career researchers, researchers from young labs, low to middle-income countries, and junior principal investigators (PIs) (Figures 1, 3, S1) (5–7). Furthermore, despite the exhausting travels to venues of these meetings, the experience of presenting at meetings for the early career researchers (ECRs) and minorities who attend has not improved appreciably (7–9). A few conferences in some disciplines have implemented valuable changes for the community and have become more receptive to attendees with families. A recent survey carried out with participants of conferences in an entire research sector, revealed that only 2% of its 2,326 respondents found these meetings to be useful and cost-effective. 44% mentioned that these conferences had “no perceptible impacts” on their research projects, programmes or policies, while 26% found conferences to be impactful, but not cost-effective (10). The increasing number and size of scientific meetings and conferences held in-person also contribute to rising atmospheric carbon dioxide (CO_2_) levels and thus to climate change, with negative implications for the research community and beyond (11,12). Scientific conferences generate multi-billion dollar expenditures, feeding business ecosystems that prey on national and international research and development budgets (13,14). The events industry describes billion dollar activity, but beyond personal value to some participants, the overall value of in-person conferences for the scientific community is seldom measurable. The status quo of academic meetings and resource allocation is not efficient, equitable or sustainable. When it comes to improving conferences, the question is not “is it possible?” but rather “are we as scientists willing to change them before we have run out of options?”. Here, we examine 270 in-person scientific conferences in various disciplines for their offered features using data available online from meeting websites. We highlight key ongoing issues and provide recommendations for improving research conferences (Figure 1, Tables S2-S11) (4). By switching from the existing in-person conference model to equitable and nearly carbon-neutral conferences, improvements will be made in a plethora of areas, with positive short- and long-term impact on research and research culture.

**Figure 1.**
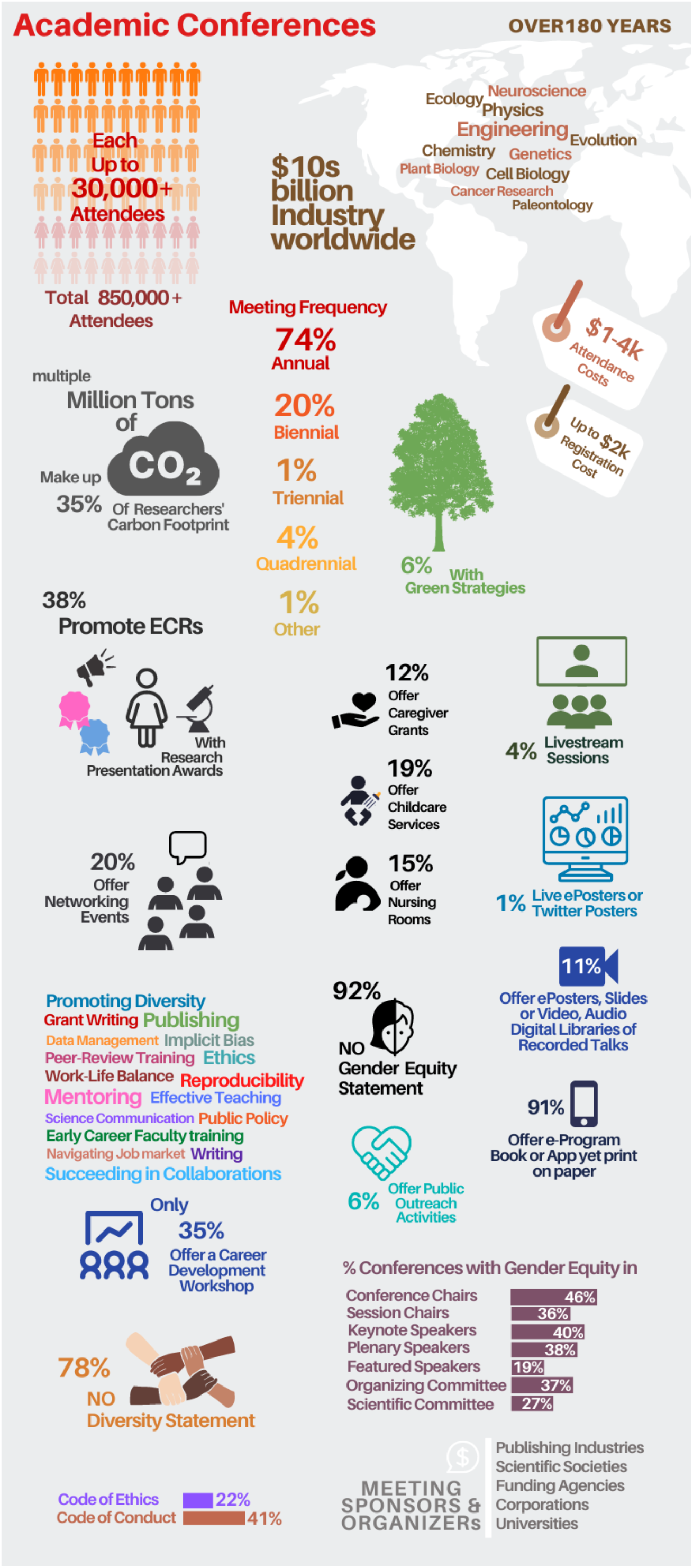
Academic conferences are costly, exclusive and environmentally unsustainable. Summary of an analysis of a database of 270 conferences held in-person and organized by over 150 scientific societies and other organizations during 2018-2019 in various scientific fields including Engineering, Physics, Mathematics, Neuroscience, Genomics, Chemistry, Ecology, Evolution, Cell Biology and Immunology (Tables S2-S11) (4). These scientific societies combined have ∼1,658,602 researchers registered members attracting a combined ∼859,114 attendees to their meetings. 1% of these meetings were held multiple times a year, 74% were held annually, 20% were held biennially and only 1% and 4% were held triennially and quadrennially respectively. Registration fees varied from US$100s to up to over $2,000 (4). Only 6% of conferences included any form of green policy. Number of attendees reached up to over 30,000 for a single event (4). A total of ∼859,114 attendees of these conferences collectively spent over US$1.288 billion of their research funds and generated over 2 million tons of CO_2_ in the form of air travel and other activities to attend these meetings (Table S6,7,8). Conference attendance represents 35%, and infrastructure 20% of the carbon footprint (15). Only 19% of all conferences offered some form of on-site childcare (free or at a cost) (Table S9). 92% of conferences did not provide a gender equity statement on members of committees and speakers and did not disclose these gender statistics on their website. 40% of conferences with information (such as first and last name of speakers and affiliations) available online had keynote speaker gender balance, 38% of conferences with information on plenary speakers and 19% of conferences with information on invited/featured speakers achieved a 50% or higher participation of women. 37% of conferences that reported information on their website achieved organizing committee gender parity and 27% achieved scientific committee gender balance. 46% of conferences reporting conference chair names and 36% of conferences reporting session chair names achieved gender parity. Only 41% and 22% of conferences examined included a code of conduct and code of research ethics or research integrity on their website respectively. The quality of these codes varied considerably among conferences. The code of conduct examined here includes both code of conduct for organizers and attendees. Only 35% included some form of a career development workshop for early career researchers (ECRs) and 38% included promotion events such as (either) special symposiums or podium talks or poster or oral presentation awards for ECRs. Only 20% of the 270 conferences offered networking events in the form of ice-breakers, mixers and meeting-with-experts sessions for ECRs (Table S9).

### The alarming carbon cost of scientific travel

The aviation industry is responsible for over 860 million metric tons of CO_2_ emissions every year, with every metric ton of CO_2_ emitted leading to 3 square meters of Arctic sea ice loss (11). With an upward of 2,500 flights a day over the north Atlantic, transatlantic flights are the third largest contributors to annual global CO_2_ emissions (Table S8). With the current career norms, researchers are expected to fly nationally and internationally, often several times a year, for scientific conferences. A single researcher’s flight from the United States to Europe and back to attend a conference will generate over 1,000 kg of CO_2_. There are 57 countries where the average person produces less CO_2_ in a year (Figures 2, S3). In addition, each night spent in a hotel creates over 70 lbs of CO_2_ from fossil fuel derived electricity. The number of attendees varies from 100 to over 30,000 for a single event (Tables S1, S6) (4), and most conferences (74%, 199/270 in our database (4)) are held annually (Table S4). This means that the average three-day, 1,000-person national conference generates over 580 tons of planet-warming CO_2_ emissions every year. The total carbon footprint from a single annual meeting of the Society for Neuroscience, hosting 31,000 attendees, is equivalent to the annual carbon footprint of 1000 medium-sized laboratories (16). The ∼859,114 attendees of 270 conferences held during 2018-2019 collectively generated ∼2 million tons of CO_2_ in the form of air travel and other activities to attend these meetings (Table S8). Multiplying these amounts of CO_2_ generated by the hundreds and thousands attending a single conference (Table S1, S8), shows why curbing emissions through diminishing scientific air travel and in-person conference attendance must become a priority for every researcher, university, scientific society, and funding agency.

**Figure 2.**
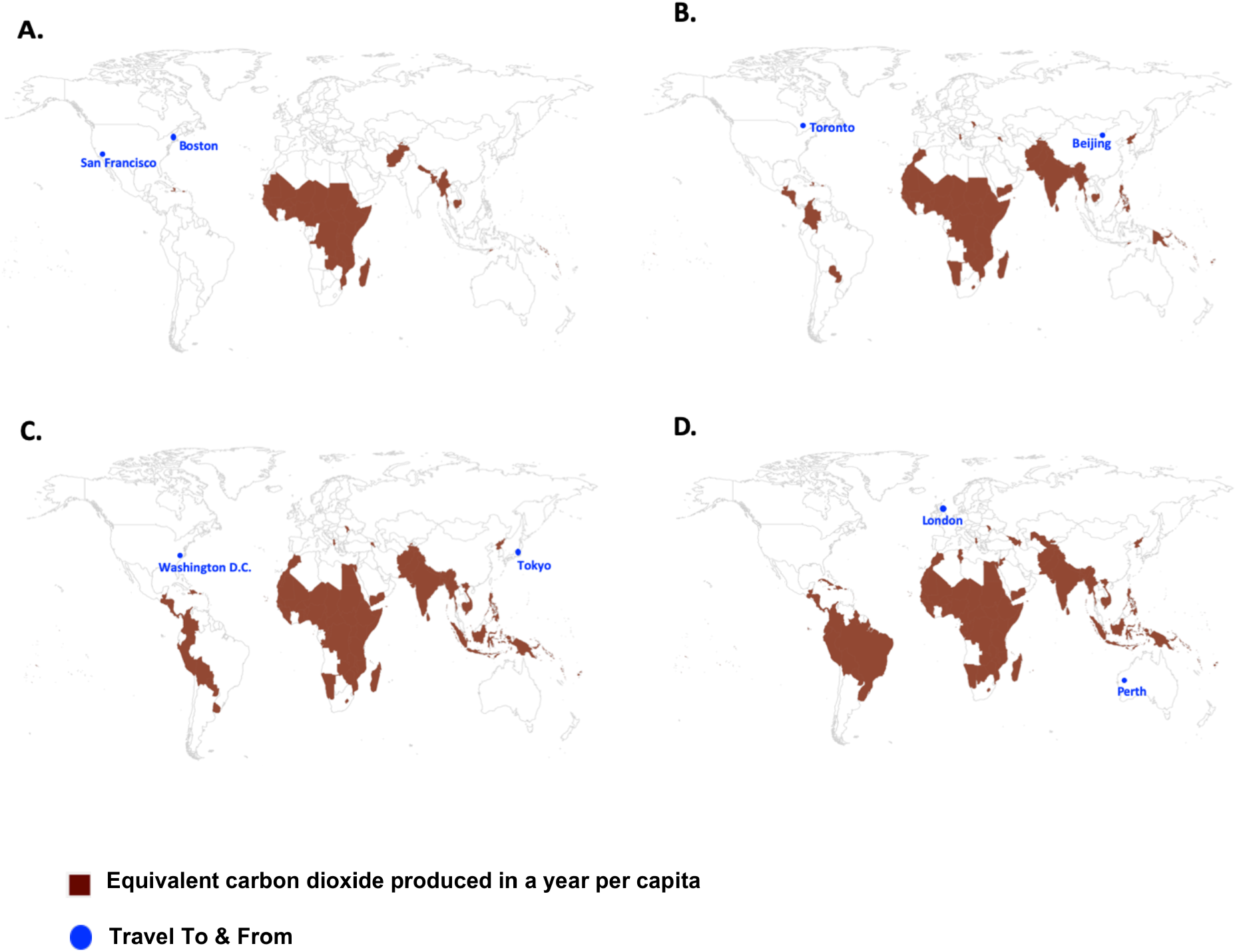
In-person conferences leave a large carbon footprint. The air travel of a *single* attendee emits as much CO_2_ as many people do in a year. Even short-haul and domestic flights produce large amounts of CO_2_. A) Flying from Boston, Massachusetts to San Francisco, California and back for an annual American Biophysical Society (BPS) conference generates about 623 kg (0.69 tons) of CO_2_. There are 47 countries where the average person produces less CO_2_ in a year (shown in color here). B) Flying from Beijing, China to Toronto, Canada and back for a Society for Neuroscience (SfN) meeting generates about 1,621 kg (1.79 tons) of CO_2_. There are 84 countries where the average person produces less CO_2_ in a year (shown in color here). C) Flying from Tokyo, Japan to Washington D.C, United States and back to attend an annual American Association for Cancer Research (AACR) conference generates about 2,144 kg (2.36 tons) of CO_2_. There are 91 countries where the average person produces less CO_2_ in a year (shown in color here). D) Flying from Perth, Australia to London, United Kingdom and back for the annual Immuno-Oncology summit generates about 3,153 kg (3.47 tons) of CO_2_. There are 109 countries where the average person produces less CO_2_ in a year (shown in color here) (Data Source, Carbon Footprint Calculator (Table S8)). Comparable to global annual per capita CO_2_ emissions in many countries (Figure S3).

Conference attendance accounts for 35% of a researcher’s footprint (15). In hosting hundreds to thousands of researchers, conferences produce substantial air travel related CO_2_ emissions (Figure 2, comparable to the global annual per capita CO_2_ emissions in many countries, Figure S3) and large amounts of other waste in the form of promotional items. The overwhelming majority of conferences are not environmentally sustainable and lack clear green strategies or climate policies (Tables S8, S9). Only 6% (15/270) of conferences we examined offered some form of sustainability initiative incorporated into their organization (Table S9). Climate change has already shown devastating consequences; extreme weather events resulting in loss of life and billions of dollars in financial costs for a single nation; the spread of viral and bacterial disease; rising sea levels; increased food insecurity; and slowing of scientific progress among many others and these are only early impacts (Table S8). Carbon dioxide emissions must drastically decrease within a decade in order to slow down the global temperature rise and irreversible climate catastrophe (Table S8). This means that time is against us and small changes in work organization and lifestyle will no longer be sufficient. There are over 8.4 million full-time equivalent researchers in the world as of 2015, representing a growth of over 20% since 2007 (Table S12). Clearly, the community that is supposed to understand the problem better should contribute these goals by reducing their travel related CO_2_ emissions.

### The cost of opportunity is prohibitively high for ECRs

The costs associated with attending conferences include meeting registration fees, airfares, accommodation, ground transportation, food and event tickets. The registration fees alone often range from a few hundred to over a thousand US dollars (Table S7). The total cost of attending large national scientific meetings in the United States is ∼US$1,000-$2,000, equivalent to one or more months of graduate and postdoctoral researcher net salary worldwide (Figure S1). As a result, a principal investigator may attend multiple conferences a year but will have to budget over ten thousand dollars to send multiple trainees to one conference per year (Table S12). The costs of attending international conferences are well in excess of national and regional meetings (∼US$2,000-$4,000). A plane ticket constitutes a substantial cost for international conferences and obtaining a travel visa is expensive (∼US$100-$1,000) (17). Trainees often have to cover the expenses upfront and on their own as travel awards are scarce and typically, only a limited number are offered by the meeting organizers, universities and non-profit organizations. In addition, these awards often barely cover the bulk of conference attendance costs, which include travel, food, visa and accommodation expenses (Table S9).

### The rich getting richer disproportionately harms ECRs

Attendance expenses may not appear burdensome to wealthy participants from select academic labs in developed nations but are unaffordable for many academic research labs worldwide. Even tenured academics can struggle to afford conference participation. The less wealthy subsidize the expenses of the speakers, who usually attend scientific meetings free of charge and benefit from these events to further build their scientific status (the Matthew effect (18)). Bursaries or reduced rates for some participants do not provide a convincing justification for why speakers, who often comprise the most well-off academics, should have their expenses paid for by other conference participants. Even if all ECRs attended free of charge, other non-keynote academic participants then may have to pay their way for the sessions. Attaining travel funds is more difficult during early career stages, and this is aggravated by funding structures and researchers’ lack of financial stability. In nations of low to middle income economies, researchers are often unable to attend national or international meetings (Figure S1) unless they are invited or manage to acquire travel grants. Furthermore, the research budget for many laboratories can be limited so that even in the rare occasions that ECRs obtain funding, they would privilege its use for research instead of traveling, as few laboratories may be able to afford both.

### Obtaining a travel visa is challenging for many researchers

Obtaining short-term visas to travel to scientific meetings is a major hurdle for many researchers, particularly those from developing nations (5). A number of countries have explicitly stated that certain nationalities are not welcome, or will have to endure lengthy, uncertain and costly procedures to obtain a visa (17). Recent long-term conflicts and changes in the political climate at conference destinations (often major scientific hubs) such as travel bans have added to these challenges. Travel bans can also come into effect during national and global health emergencies such as epidemics and pandemics. These obstacles not only deeply affect the lives and careers of scientists working in developing countries, but also citizens of countries with visa restrictions who live and perform research in global scientific research centres planning to attend conferences elsewhere (Figures 3, S2).

**Figure 3.**
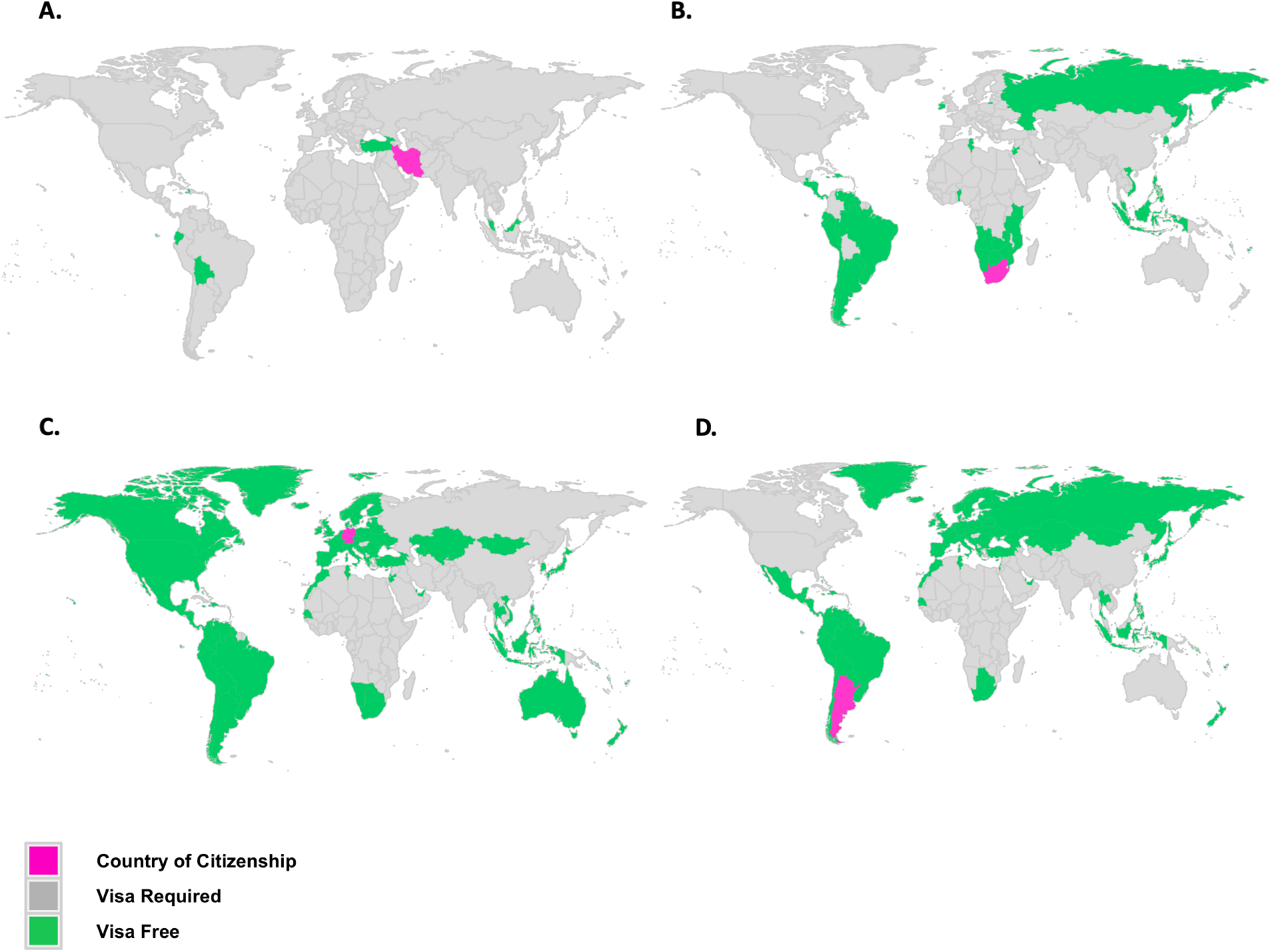
Visa restrictions limit scientific career development. Short-term visitor visa requirements for researchers who are citizens of A) Iran B) South Africa C) Argentina and D) Germany. Researchers who are citizens of these four countries (in magenta) can only travel to select countries (in green) without applying for a visa for a short visit to attend a scientific conference (see more in Figure S2).

### Conferences need to improve on equity & inclusivity

Equity in planning scientific meetings creates fair opportunities for all and inclusivity creates a welcoming environment for all participants. The current organization of many conferences leads to practices that exclude researchers on a wide range of factors including, but not limited to, gender, ethnic, racial, socioeconomic, health and geographical backgrounds, and career stage. For instance, women and researchers from racial and ethnic groups, who are under-represented in various fields, are the least likely to be offered opportunities to speak at or chair meetings in their discipline (7,19). 92% of 270 conferences that we examined lacked a statement of speaker gender balance and 78% lacked a diversity statement (Figure 1, Table S9). At 270 conferences in various disciplines which reported names of chairs, organizers and invited speakers online, women were underrepresented among conference and symposium session chairs, plenary and keynote speakers, invited lecturers, or as panelists in a broad range of academic meetings and disciplines (Figure 1, Tables S9, S12). Only 46% and 36% of conferences achieved gender parity for conference chairs and session chairs respectively; 37% and 27% achieved gender parity for conference organizers or steering committees and scientific committee respectively; 40% and 38% achieved gender parity for keynote and plenary speakers respectively and only 19% had equal numbers of male and female featured speakers (Table S9, Figure 1). The process by which speakers are selected, including speakers chosen from abstract submissions, can generally be improved by removing intended and unintended biases (20). The current lack of diversity hinders advancement in scholarship, and it is especially discouraging to junior researchers (Table S12). There has also been inertia in setting up and implementing more family-friendly policies at scientific meetings. Free and on-site lactation rooms and childcare was only provided at 15% and 19% of meetings respectively, while caregiver grants were only offered at 12% of conferences examined (Table S9, Figure 1) (8). In addition, many of the currently offered lactation rooms do not have the minimal tools to provide an adequate environment. Only 35% of conferences included some form of a career development workshop for ECRs and 38% included promotion events such as (either) special symposiums or podium talks or poster or oral presentation awards for ECRs only (Tables S9, S10, S11). 20% of the 270 conferences offered networking events in the form of ice-breakers, mixers and meeting-with-experts sessions for ECRs (Table S9, S11).

### Improvements to academic conferences

#### Replace in-person national and international meetings with more sustainable travel to regional meetings

Researchers should be at the forefront of reducing CO_2_ emissions. This can also impact the wider scientific community regarding changes in academic practices and alter public opinion regarding changes in travel and lifestyle (21). Conferences in far-off vacation destinations are difficult to justify as these accumulate a large carbon footprint, often without clear scientific objectives and little to no benefit to the local scientific community. Most ground-based transportation produces lower CO_2_ emissions (Table S13, example 2). Research institutions, funders, and scientific societies also need to incentivize and mandate regional meetings (albeit less frequently) (Table S13, example 5) in place of large national and international gatherings. Smaller regional event sizes enable attendees to interact and discuss science in ways that provide ECRs with ample opportunities to exchange ideas and meet established researchers. Regional society meetings provide many benefits, such as low hosting costs. These can also be live-streamed, recorded and made available online for the benefit of other researchers globally. Regional conferences held in more economic and public venues, schools or university campuses, may also offer attendees the opportunity to visit local laboratories, tour facilities, and interact with the local scientists, potentially bringing benefit to the local community (Box 1).

#### Make research results more accessible globally in virtual mode

Face-to-face interactions at conferences that are important in generating new ideas, networks and collaborations can also be facilitated via digital conferencing (Table S13, example 1). Reducing academic air travel does not impact professional success (22). Speakers spend time and resources preparing talks for in-person conferences which end up reaching a limited audience with limited geographic impact that is often restricted to wealthy economies. Switching to fully virtual conference modes will increase reach and access to knowledge worldwide. Multi-location in-person conferences where participants only travel to a nearby location (conference hub broadcasted digitally) to interact with other “local” scientists are also viable (23). These semi-virtual conferences can be distributed across multiple continents and linked live digitally (24). Each hub may hold in-person keynote and panel sessions, but the hubs can also be connected with each other for virtual panel sessions and discussions; such an arrangement enables speakers to present talks and posters remotely. Live streaming allows chat amongst other online attendees in real time, access to sessions that fit attendee schedules, and the ability to stream sessions from any devices while on-the-go creating more space for researchers to interact and connect. The number of conferences that are live-streamed or held in a virtual reality setting have significantly increased over the past few years (Table S13, example 6), particularly the last months in the context of the COVID-19 outbreak (Table S13, example 7) (25), showing great success (Figure 4) (26,27,66). Nonetheless, many more are needed in all disciplines.

**Figure 4.**
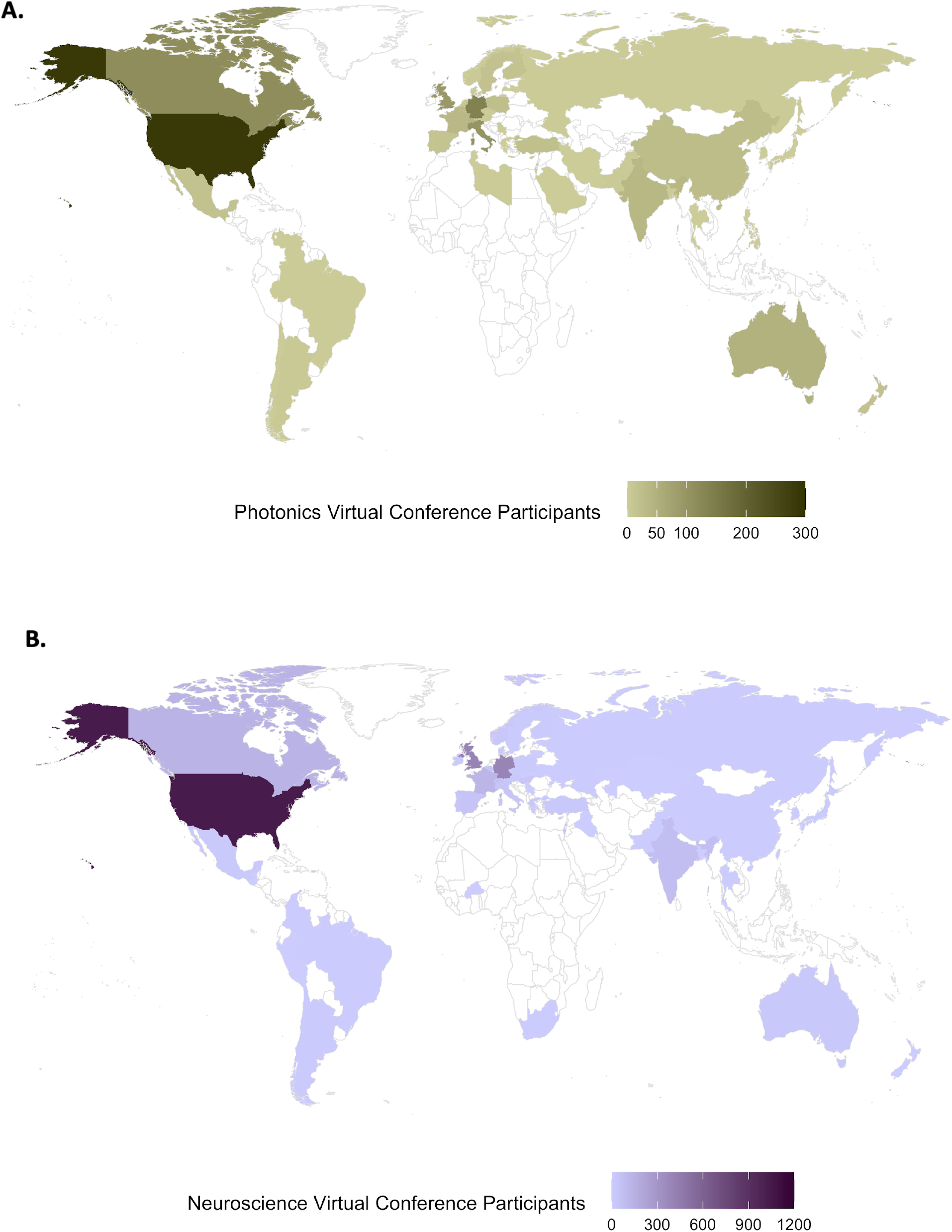
Virtual conferences attract wide global attendance. A) The Photonics Online Meetup (POM) held on January 13, 2020 as a virtual conference with over 1,149 students and researchers interested in optics gathered to participate in the as groups, creating local communities from 63 research hubs, 47 countries on 6 continents. The meeting had three themes, submitted abstracts were chosen for either presentations or posters and the presentations were held in real-time (26). B) The Neuromatch virtual conference was held on March 30-31, 2020 attracted a total of 2,930 attendees from 56 countries (27). With over 70% of attendees comprising graduate and postdoctoral trainees, Neuromatch 2020 meeting included poster sessions, short talks, invited and contributed talks as well as one-on-one meetings. Neuromatch 2020 also achieved an equal number of men and women in invited speakers (27) (Raw Data on attendee numbers and geographic locations courtesy of Drs. Andrea Armani, University of Southern California and Konrad Kording, University of Pennsylvania).

#### Foster digital networking by investing in relevant, immersive and interactive experiences

It is crucial for organizers of virtual conferences to develop strategies for facilitating digital connections. Fully streaming conferences (beyond video-conferencing select talks) and scientific electronic panels can be made feasible without many significant additional resources, accompanied by online discussions and Twitter (poster) conferences (28,29) (Table S13, examples 1,8). Participants can also record and upload videos of their presentations that are posted to the meeting website. During several weeks when the conference is “open”, attendees can watch the videos, ask questions, and interact with the speakers on online forums; educational and hiring opportunities can be offered and applied for. Slack groups have taken the initiative to connect researchers in life sciences related disciplines globally via group based and one-on-one online discussions and keep them virtually connected afterwards (30,31). Electronic (online) poster sessions can provide each presenter with the opportunity to tag their poster with a 3-5 minute talk summarising their work. Attendees watching video presentations of talks and posters can ‘direct message’ the authors/presenters for inquiries via email or the online meeting platform. Some conferences cancelled due to the COVID-19 pandemic launched their meetings online using *Open Science Framework (OSF)* Meetings to facilitate virtual dissemination of presentations and posters. Digital conferences and discussion forums can serve to assist communication between early career and senior researchers since writing a comment or question in a forum can feel less intimidating than approaching an established scholar in person. With more time to think the presentations through, attendees can put their best foot forward during interactions and networking with speakers.

#### Preprinting research outputs could reduce the number of all conferences and can improve digital conferences

There is a need for researchers to critically evaluate the necessity of holding conferences. Early dissemination of scientific findings can occur online via preprints. Preprints are written scientific outputs uploaded by researchers to public servers such as *arXiv, bioRxiv* and *OSFramework. BioRxiv* and *medRxiv* currently host in excess of 100,000 manuscripts combined, receiving over 4 and 10 million views per month respectively (32). Preprints are already benefiting life scientists at large, but can be used in new ways to aid career development and increase the efficiency of scientific research and communication. Many investigators, especially in countries of low to middle economies, are now able to disseminate and read the most recent updates in their field via preprints months to years prior to reading new results in journals, further reducing the need to travel to discuss already published findings. Additionally, all conference materials such as abstracts, videos of talks, posters and presentation slides can be uploaded to open access platforms such as *Zenodo, protocols*.*io*. or channels dedicated to preprinting conference proceedings on *bioRxiv* with a Digital Object Identification (DOI). Such sharing does not entail extra work for researchers, enables citation of these outputs on manuscripts and curriculum vitae and allows review on-site or via peer-review platforms such as PREreview (33). Digital conference libraries allow researchers in all time zones and those unable to attend these conferences asynchronously. Widespread preprinting of research results can reduce the number of conferences held overall (Table S13, example 14).

**Box 1.** Actions to Improve small & large, hybrid & fully virtual conferences.

| **Box 1. Actions to Improve small & large, hybrid & fully virtual conferences** |  |
| --- | --- |
| **Replace in-person national & international conferences with fully virtual & regional meetings** | Hold small and large meetings fully online or connect regional in-person conference hubs digitally by live streaming the conference. Large conferences are often hosted in expensive cities as there are many accommodation options for large crowds, while conferences in more affordable locations are typically smaller in size. Use online platforms for your virtual academic meetings (Table S13, example 1). Examples: *EBRS 2019 carbon reduced meeting*, *Neuromatch 2020* and *POM 2020 meeting*. |
| **Increase accessibility for researchers with audio-visual difficulties** | Provide live closed captions and subtitles (are offered by leading video conferencing systems) to help individuals with impaired hearing and those who are not fluent in the main language of the conference, or have missed certain parts of the presentation (57). Offer transcripts to attendees after the talks. Instruct the presenters to cover the whole content of every slide, describe all images, and include an audio described version of any video that is played, to become more inclusive to attendees who are visually impaired. |
| **Assist attendees with internet connectivity or financial challenges** | Provide internet hotspots at low cost to enable participation. Attendees in remote geographic locations nationally may not have access to high-speed internet to participate in virtual meetings. Help attendees who are unable to afford the conference registration fee, to source local sponsors for support. |
| **Go Paperless: Stop printing meeting programs and announcements** | Printing program books of 200 pages each for 15,000 conference attendees uses 3 million pages of paper (~150-300 trees). Instead provided complete electronic program schedules accessible electronically using mobile phone applications online or offline saving up to a few hundred thousand to millions of pages of paper at a single large conference. Paper should be used minimally, if at all, at conferences and any paper used (e.g. for promotional material or signs) should be 100 percent recycled. Examples: *Evolution 2020* meeting. |
| **Eliminate conference merchandise & non-compostable products** | Conference badges, bags and various other paraphernalia create large wastage and cost organizers and sponsors significant sums of money. The cost of ordering souvenirs can be repurposed to create more ECR promotion events and training workshops. Ask attendees to bring/reuse their own materials and take notes electronically on their portable devices. Use seed paper conference name badges (Table S13, example 4). Lanyards and gift mugs can be made from re-usable or compostable material or excluded altogether. Examples: *Evolution 2020* & *I Scientist 2019* meetings. |
| **Select environmentally responsible accommodation** | For in-person regional meetings, designate attendees to accommodations that operate in an environmentally responsible manner. Select hotels at locations near public transportation. Examples: *Evolution 2020* meeting. |
| **Select sustainable food catering** | Conference dining menu can improve by serving less meat (34), particularly beef and pork products, and increasing relative representation of quality vegetarian and vegan options, prepared with locally sourced, in season ingredients at the catered events. Examples: *Evolution 2020* & *ISHPSSB 2019*. |
| **Reduce plastic usage** | Disposable cups should not be provided during refreshment breaks. Attendees can bring their own reusable water bottle (and mug) and conference organizers can make several hydration points available. Waste can also be reduced by using compostable food service materials whenever possible (34). Organizers should make an effort to push for reusable crockery via access to a kitchen, non-disposable (made of ceramic, washed and reused) dishware and cutlery when serving food is included in the program. Examples: *Evolution 2020*, *Gordon Research Conferences* & *I Scientist 2019 meeting*. |
| **Include environmental improvement events** | Arrange for environmental improvement events such as clean-up as a social activity in the form of forest or beach walking clean up that can also serve as photo opportunities for conference social media, providing time for attendees to socialize and discuss science while helping the environment, simultaneously achieving multiple goals. |
| **Organize public outreach activities for attendees** | Few conferences include public outreach events (e.g. public lectures). A possible social activity that could build longer-lasting connections is reaching out to local schools and libraries so that researchers attending the conference can also give public lectures with a layperson version of their talk or poster and engage with the public. Examples: *Ecology & Behaviour Conference 2019*. |
| **Provide free & on-site nursing & childcare facilities at in-person meetings** | At most conferences, free or on-site facilities to support special needs such as lactation rooms for nursing mothers, childcare and assistance for parents or other caregivers of children and children's playrooms, are neither available, affordable, nor designed for female users. Lack of support and the complexities of arranging care can make travelling for conferences extra expensive and stressful for ECR parents who must choose to either miss out on the career opportunities of travel, or incur costs and significant personal stress to them and their family. Providing such utilities is completely feasible and well within the interests of conference organizers, funders, scientific societies and attendees (35,36). Examples: *Evolution 2020*, *I Scientist meeting* meetings (Tables S13, examples 19,20). |
| **Generate meeting code of conduct** | Majority of in-person conferences lack a code of conduct (37,38). Larger and national conferences are more likely to have codes than international or smaller ones - emphasizing the need for increased efforts in order to improve inclusivity and diversity and reduce the loss of (historically) marginalized scientists. Scientific societies, funding agencies and conference organizers should publish a code of conduct upfront for all participants to sign during conference registration and abstract submission processes and strictly enforce the policy. Examples: *R conference*, *Molecular Biosystems Conference* (Tables S13, example 17). |
| **Generate meeting code of ethics** | Most conferences lacked a code of research ethics for attendees on the meeting or associated scientific society websites. Include a code of research integrity for every meeting. Examples: *Society for Ethnobiology meeting 2019*, *Society for American Archaeology meeting* (Tables S13, example 18). |
| **Improve attendee physical safety** | Not all conference locations are safe for attendees, especially for women (39) and few provide safety instructions for attendees. Organizers could implement mandatory safety apps (40), a buddy system where attendees accompany or warn and protect each other and report any issues that jeopardize their safety. Meeting organizers must also choose safe accommodation locations, safe and affordable transportation modes, less emphasis on alcohol focused events and put forward anti-harassment policies and establish a clear system for reporting and handling such issues if they arise. Smaller and regional conferences can enable attendees to look out for one another. Examples: *Evolution 2020 meeting* (Tables S13). |
| **Organize well-thought networking activities** | Well-planned networking events (e.g. ice-breakers) are necessary to introduce and to connect researchers across career stages within and across disciplines (without alcohol and with a focus on the event). Researchers can be randomly seated together or assigned to breakout rooms during virtual conferences and tasked with coming up with an idea for a collaboration in a project planning workshop or seek mentorship. Examples: The *EMBO/EMBL "Seeing is Believing" 2019 meetings* (Table S13). |
| **Create a central database for scientific conferences** | Conferences are often not well-characterized, evaluated or promoted within and across disciplines and countries; making it difficult and time-consuming for researchers worldwide, in particular ECRs, to identify the most beneficial (Figures 5, 6) gatherings. Attendee and speaker gender and career stage statistics are seldom reported online by conference organizers. A central database (4) can help keep researchers well-informed on the past and future meetings and the key considerations of environmental sustainability and equity that all scientific conferences must possess (4). |

#### Incentives and mandates to support and promote low-carbon research careers

Business-related air travel by academics at universities continues to grow, reaching tens of millions of miles flown by their academics in one year (41,42). Institutional practices should shift to mandate severe reductions in academic air travel as these changes will significantly reduce the academic carbon footprint (16) (Table S13, example 3,9,11). As researchers re-evaluate their practices, it is essential that academic institutions, scientific societies and funding agencies coordinate and mandate sustainability code of conduct and funding to only organize academic conferences in ways that require less flying (43). At many institutions and disciplines, scholars are granted tenure and promotions based in part on the number of their research presentations at professional conferences (44,45). Furthermore, as postdoctoral and tenure-track jobs are scarce in many academic disciplines, ECRs face mounting pressure to attend conferences in order to network with potential colleagues. Changes can also be expected in funding and promotion requirements so that tenure-track academics do not have to choose between delivering an international talk and receiving a grant or tenure promotion. The proposed changes will in turn encourage a culture of academic research that is less reliant on traveling. Some research institutions and funders have implemented a carbon tax on air travel emissions caused by academic flying; other institutions have made it compulsory to travel by train if the travel time is short. Funds from a carbon tax on ground travel can also be used to fund research that could fight against the climate crisis (42,46). Although institutions can estimate total emissions and buy offsets (by paying for renewable energy or programs designed to reduce emissions), the damaging impact of researchers flying on climate and living ecosystems are irreversible and the scale of the efforts and cost to actually offset emissions from flying may be prohibitively high (47,48). A shift towards regional and digital conferences would be more effective and environmentally sustainable.

#### Improving intersectionality and career stage equity at regional and digital conferences

Conscious efforts must be made by the organizers of every conference to maintain fair representation of all researchers (9,49,50). Few conferences publish a statement of gender equity, diversity or inclusion and actually achieve inclusion of 50% or higher female speakers (Table S13, example 16). Maintaining gender balance impacts both speakers and ECRs, who need to see representation and role models of their own gender in their field. Without clear guidelines and targets, unreasonable excuses are made for lack of diversity. Claims that, “there are no available female speakers or not enough women in a field” are no longer acceptable, given recent increases of minority researchers in many scientific fields across graduate, postdoctoral and faculty career stages (51). Oftentimes not all avenues for seeking female speakers are not explored (52). Overcoming biases involves not only awareness but also positive action. Blinding the selection of talks, removing conflicts of interest, utilizing open databases listing ECRs from underrepresented groups (53), and having fewer long talks to make time for more short talks may all help ensure that minority groups of scientists are welcomed and equally represented in conferences (20,54) (Table S13, example 10). Funders and meeting organizers need to pledge and enforce gender equity policies and public inclusion declarations and guidelines, and publish the names of invited speakers and session chairs in advance of the conference program. Conference attendees can express conditional attendance, lifted only after the invitation of men and women speakers in a fair and balanced manner. In some disciplines, such as Ecology, multiple (but not all) conferences are yielding towards achieving gender equality for speakers and chairs; aiming beyond existing representation at the faculty level. The number of ECRs has increased globally, and a number of talks can be allocated to early and mid-career researchers. Senior faculty can nominate an ECR trainee or recommend an ECR faculty in their place to attend the conference. Similar guidelines can be adopted for the selection of session chairs and audience members volunteering to ask questions after each talk. A gender disparity in participation during discussions exists as discussions are most often led by professors (13,14). To change this, every session can be chaired by at least one ECR and have at least one ECR organizing the Q&A discussions. Trainees may also not volunteer to ask a question, which can be a daunting task. A good policy can be to specifically open the session to trainees, or, if the talk is streamed online, to take the first question from the internet using platforms that allow audience interactions (Table S13, example 1).

#### Improving trainee and ECR faculty career development

Conferences are an optimal setting for diversity events, academic and non-academic job search training, training in strengthening collaborations, scientific communication, scientific writing, reproducibility, career development teaching, mentoring workshops and more. Yet a minority of recently held academic conferences offered some form of a career development workshop (4). The “Reproducibility for Everyone” group of the *eLife* community ambassadors has pioneered reproducibility workshops at a number of conferences (55). There are few, if any, advocacy events such as featured talks and poster sessions for up-and-coming ECRs, for instance, those going on the academic or non-academic job market or junior faculty. All conferences can implement ECR promotion events in the form of special symposia or awards. These would bring many benefits to women and other underrepresented groups, which typically lack support, sponsorship and advocacy, and thus miss out on career opportunities.

### Benefits of the proposed improvements to Research and Research Culture

#### Digital conferences are environmentally sustainable

In-person conference attendance generates multiple million tons of CO_2_ (16,56). Only 5% of the global population enjoy the privilege of flying annually and this includes a subset of academics. A number of scholars routinely fly over 100,000 miles every year. Academic air travel makes up almost half of total university emissions (Table S13, example 9). Other damaging aspects of conferences include plastic badges, bottles, paper and other promotional items and unsustainable food catering to masses of attendees (15) (Box 1) (Table S13, example 15). Reducing travel to national and international conferences and organizing virtual conferences will have a dramatic impact on the environment. For example, a recent experience with multi-site conferencing showed reductions of 50%-70% in travel-related greenhouse gas emissions compared to the single-site alternatives despite increased participant numbers (23). Low-cost or free nearly carbon neutral conference models (Figures 4, 5, 6) will be of great benefit to both our scientific communities and our planet (57).

**Figure 5.**
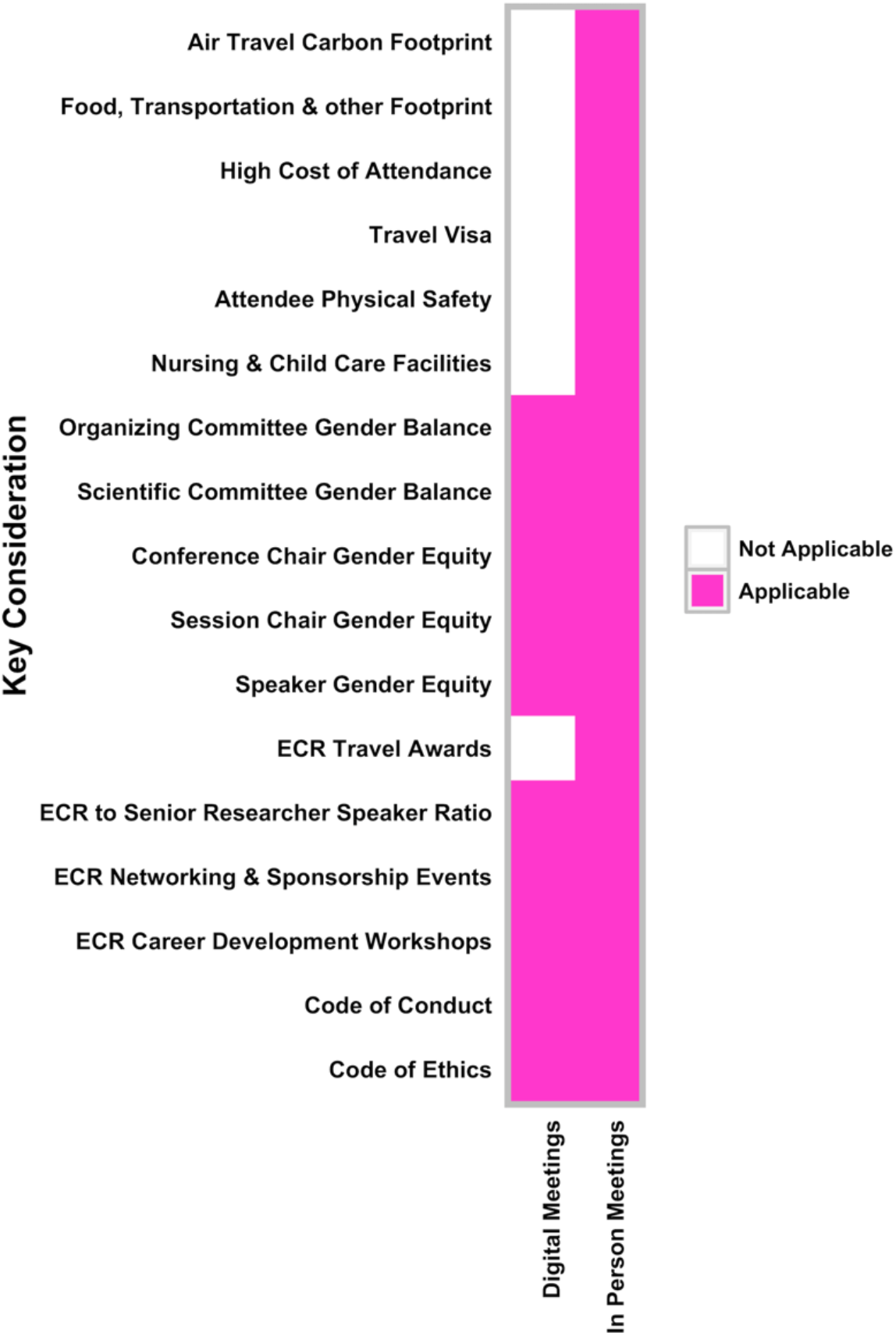
Digital scientific meetings present many advantages for the diverse scientific community. A visa (travel permit), travel carbon footprint, early career researcher (ECR) travel awards, in-person attendance costs such as flying, accommodation, transportation and meals, childcare and infant nursing facilities and attendee physical safety are not applicable considerations to digital (online) conferences. Virtual conferences do not require catering, venue rental, or on-the-ground logistics coordination, thus are substantially less expensive to organize than fly-in conferences.

**Figure 6.**
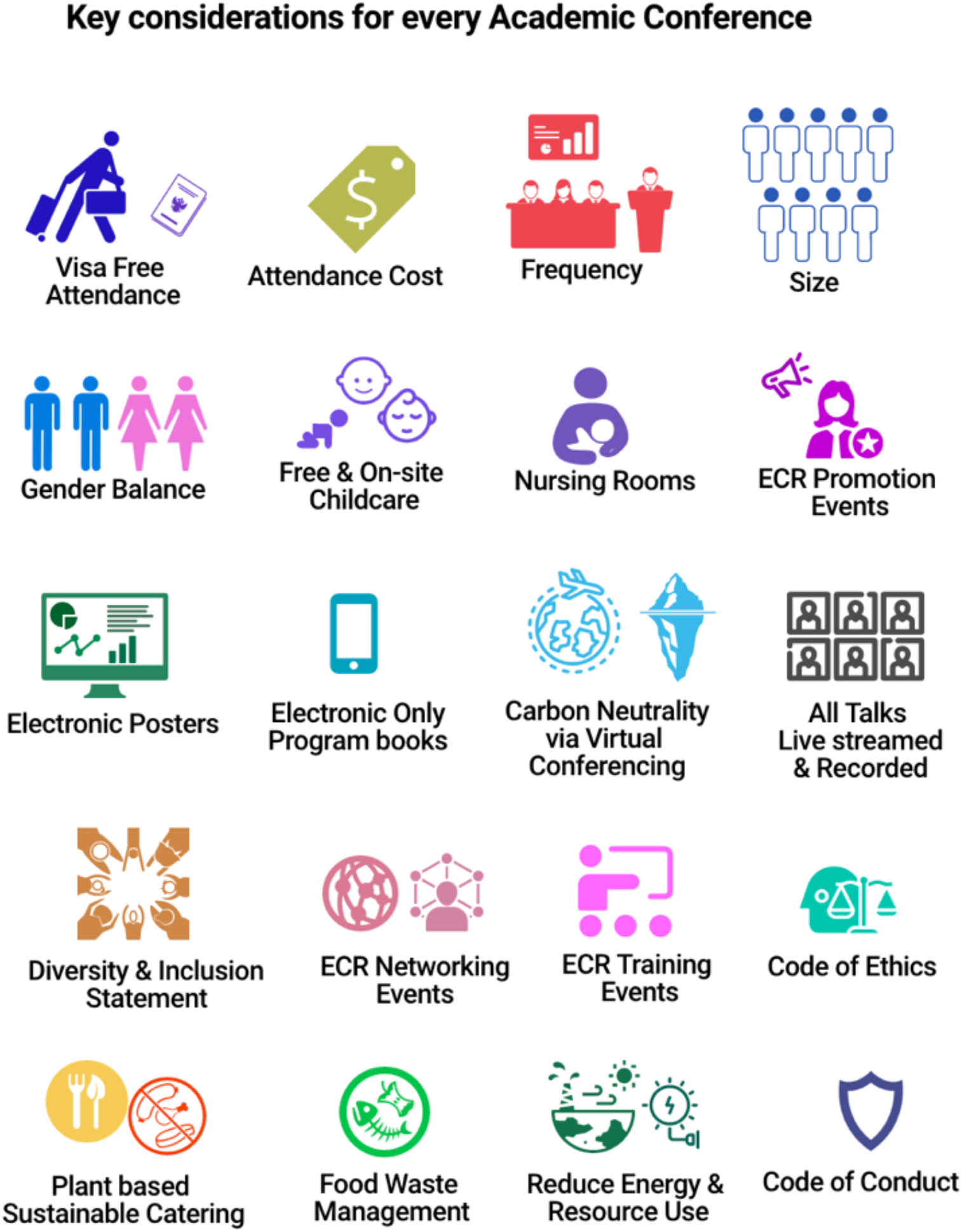
Key considerations for every Academic Conference. A checklist of key considerations for in-person or virtual regional conferences, virtual national, and international conferences for researchers, funders, scientific societies and conference organizers. ECR: Early Career Researcher.

#### Digital conferences return funds to researchers

The global market size for all academic meetings and events is estimated at US$11.5 billion per year, growing annually (Table S13, example 12) (13,58). A total of ∼859,114 attendees of 270 conferences, collectively spent over US$1.288 billion during 2018-2019 to attend these meetings (with a total attendance cost average at US$1,500 per attendee). This is over 3.3% of the NIH total annual budget. Conference organizing has become an industry, generating income for commercial or society conference organizers; travel, hotel and catering industries (14); and local tourist attractions that have little interest in inclusivity or the academic content being discussed. Researchers who attend expect many benefits, such as diverse early career researcher training sessions or finding their next collaborator, but most conferences have never been evaluated for their actual impact on their field. A fraction of research funds used to cover attendance of in-person conferences can be devoted to establishing digital facilities at universities and other research institutions so that scientists globally can attend meetings online. These resources do not require large funds and will be of benefit to the researchers and institutions that invest in them, long after the conferences have ended.

#### Digital conferences are more inclusive

Fly-in conferences may meet the needs of some academics and professionals in wealthy countries but researchers with family commitments, physical or health limitations and vulnerabilities cannot easily attend these conferences. Digital conferences incur lowest costs and can be organized when travel is not possible due to disease outbreaks such as the recent COVID-19 pandemic (25) (Table S13, example 13). Graduate and postdoctoral trainees on average produce about three times less emissions from air travel than fully tenured professors, and female academics also travel less than their male colleagues (59,60). Virtual meetings can boost the number of registered participants up to 50% compared to in-person conferences (61,66). In addition to the added socio-economic and ethnic diversity, digital conferencing can increase diversity of the career stages and genders of attendees, which in turn improves the quality of the scientific research and the conferences (51,52,54,62). Virtual conferences also present an upper hand regarding accessibility issues. Live closed captions and subtitles (offered by leading video conferencing systems) may not only be a way forward for individuals with impaired hearing, but also for those who are not fluent in the main language of the conference, or those who missed certain parts of the presentations (63). Delegates may also receive a transcript of the presentations afterwards. Digital conferences can also become inclusive for visually impaired individuals, by utilising text-reader friendly documents and by instructing presenters to cover the whole content of every slide, describe all images, and include an audio described version of any video that is played. Software developers and organizers will have to invest more efforts into exploring the best approach to enable these features virtually.

## Conclusions

We examined 270 in-person conferences for their environmental impact, inclusivity and accessibility features. Our analysis shows that a growing body of researchers from diverse nationalities and backgrounds face difficulties in obtaining means to attend in-person conferences. Our analysis also shows that there is considerable scope for improvement in the organization of these meetings. Technological advances provide a unique and timely opportunity to reform scientific conferences. Multi-hub hybrid and fully virtual conferences can dramatically improve the experience for all. Widespread digitization will increase global access to knowledge during and after conferences, by increasing the diversity of speakers who are restricted in travel and will also enable greater democratisation of science – equalising for differences between researchers of lower funding and prominence. One can imagine a new reality where streamed and recorded conference talks will become a new integral component of the global knowledge economy. Broader access to the latest scientific developments will further accelerate knowledge gain and sharing of ideas and innovation everywhere, leading to better use of researchers’ time and funds, which will be more beneficial for science and the broader public. Digital conferences further enable scientific communities to substantially reduce the environmental impact of these meetings, meaningfully contributing to the global goal of reducing greenhouse gas emissions. Eliminating the large in-person national and international conference bubble and their costs will return funds to research labs and enable more trainees and underrepresented groups to attend and exchange their ideas. Accessibility of conferences for researchers with disabilities needs to be examined in future studies, and should be kept in mind even when switching to virtual conference formats. It will be necessary to radically reassess the exact purpose of scientific meetings and to recraft the conference format so that diverse researchers can attend and build relationships that are essential for scientific success. Addressing these challenges needs a will from scientific societies, in order for these gatherings to be truly international. Science aspires to contribute to the collective knowledge, solutions of humanity’s greatest challenges and betterment of the human condition. Therefore, they must maintain consistency and responsibility to create conditions that allow science to thrive, both socially and environmentally. The prevalence and continuity of the issues associated with in-person scientific conferences call for all researchers, their institutions, scientific societies, and funders to take initiative to recognize the problems, demand change, and actively participate in reorganizing conferences to create more open, inclusive, and kinder research environments as we advance knowledge.

## Supporting information

Supplementary File_Sarabipour et al_2021

## Acknowledgements

The authors would like to thank Drs. Richard Sever, Ronald Vale, Alexandre Bisson Filho, Dominic Trépel, Edward Emmott, Renuka Kudva, Steven Burgess, Jason Fernandes, Andy Tay, Vinodh Ilangovan and Inez Lam for valuable comments on an earlier version of the manuscript.

## References

1. Cyranoski D, Gilbert N, Ledford H, Nayar A, Yahia M. Education: The PhD factory. Nature. 472(7343):276–9 (2011).

2. Statistics Regarding International Conferences held in Japan in 2017. Japan Meetings & Events. Available from: https://www.japanmeetings.org/plan-your-event/statistics/statistics.html

3. Lindsay R. Pool RMW, Lindsey L. Scott DR, Rediet Berhane CW, Katrina Pearson JAS, Walter T. Schaffer. Size and characteristics of the biomedical research workforce associated with U.S. National Institutes of Health extramural grants. FASEB J. 30:1023–36 (2016). Available from: https://www.fasebj.org/doi/full/10.1096/fj.14-264358?url_ver=Z39.88-2003&rfr_id=ori%3Arid%3Acrossref.org&rfr_dat=cr_pub%3Dpubmed

4. Improving Conferences: A database for scientific conferences and their key features. Available from: https://elifeambassadors.github.io/improving-conferences/

5. Waruru M. African and Asian researchers are hampered by visa problems. Nature (2018); Available from: https://www.nature.com/articles/d41586-018-06750-1

6. Melonie Fullick. It’s time to rethink academic conference funding. University Affairs. (2016). Available from: https://www.universityaffairs.ca/opinion/speculative-diction/its-time-to-re-think-academic-conference-funding/

7. Ford HL, Brick C, Azmitia M, Blaufuss K, Dekens P. Women from some underrepresented minorities are given too few talks at world’s largest Earth-science conference. Nature 576:32–5 (2019). Available from: https://www.nature.com/articles/d41586-019-03688-w

8. Langin K. Are conferences providing enough child care support? We decided to find out. Science (2018). Available from: https://www.sciencemag.org/careers/2018/12/are-conferences-providing-enough-child-care-support-we-decided-find-out

9. Larson AR, Sharkey KM, Poorman JA, Kan CK, Moeschler SM, Chandrabose R, Marquez CM, Dodge DG, Silver JK, Nazarian RM. Representation of Women Among Invited Speakers at Medical Specialty Conferences. J Womens Health. 29, 1–12 (2019) Available from: https://www.liebertpub.com/doi/full/10.1089/jwh.2019.7723?url_ver=Z39.88-2003&rfr_id=ori%3Arid%3Acrossref.org&rfr_dat=cr_pub%3Dpubmed&

10. Biswas AK, Tortajada C. Impacts of Megaconferences on the Water Sector. 1st ed. Springer-Verlag Berlin Heidelberg; 2009. XVI, 276. Available from: https://www.springer.com/la/book/9783540372233

11. Notz D, Stroeve. Observed Arctic sea-ice loss directly follows anthropogenic CO2 emission. Science 354(6313):747–50 (2016). Available from: https://science.sciencemag.org/content/354/6313/747

12. Joseph Nevins. Academic Jet-Setting in a Time of Climate Destabilization: Ecological Privilege and Professional Geographic Travel. Prof Jeographer. 2013 May 2;66(2):298–310. Available from: https://www.tandfonline.com/doi/full/10.1080/00330124.2013.784954

13. Rowe N. The Value, Scope and Cost of Conferences: looking beyond the Events industry. In. p. Symposium 5788. (2017) Available from: https://www.srhe.ac.uk/conference2017/abstracts/0068.pdf

14. Row NE. The Economic Cost of Attending Educational Conferences. Int J Soc Educ Sci.1(1):30–42 (2019).

15. Achten WMJ, Almeida J, Muys B. Carbon footprint of science: More than flying. Ecol Indic. 34:352–5 (2013). Available from: https://www.sciencedirect.com/science/article/pii/S1470160X13002306

16. Nathans J, Sterling P. Point of View: How scientists can reduce their carbon footprint. eLife. 5, e15928 (2016). Available from: https://elifesciences.org/articles/15928

17. Gewin V. What scientists should know about visa hurdles. 569:297–9 (2019). Available from: https://www.nature.com/articles/d41586-019-01428-8

18. Bol T, de Vaan M, Van de Rijt A. The Matthew effect in science funding. Proc Natl Acad Sci USA, 115(19):4887–90 (2018). Available from: https://www.pnas.org/content/115/19/4887

19. Else H. How to banish manels and manferences from scientific meetings. Nature 573:184–6 (2019). Available from: https://www.nature.com/articles/d41586-019-02658-6

20. Vallence AM, Hinder MR, Fujiyama H. Data-driven selection of conference speakers based on scientific impact to achieve gender parity. PLoS One. 4(7):e0220481 (2019). Available from: https://journals.plos.org/plosone/article?id=10.1371/journal.pone.0220481

21. Reducing emissions from aviation. European Union: European Commission Climate Action; (2019). Available from: https://ec.europa.eu/clima/policies/transport/aviation_en

22. Wynes S, Donner SD, Steuart Tannason NN. Academic air travel has a limited influence on professional success. J Clean Prod. 226:959–67(2019). Available from: https://www.sciencedirect.com/science/article/pii/S0959652619311862

23. Coroama VC, Hilty LM, Birtel M. Effects of Internet-based multiple-site conferences on greenhouse gas emissions. Telemat Inform. 29(4):362–74 (2011). Available from: https://www.sciencedirect.com/science/article/pii/S0736585311000773

24. Abbott A. Low-carbon, virtual science conference tries to recreate social buzz. Nat News (2019); Available from: https://www.nature.com/articles/d41586-019-03899-1

25. Biogen’s Boston conference continues to be a leading source of coronavirus cases. Boston Herald (2020). Available from: https://www.bostonherald.com/2020/03/09/biogens-boston-conference-continues-to-be-leading-source-of-coronavirus-cases/

26. Reshef O, Aharonovich I, Armani AM, Gigan S, Grange R, Kats MA, et al. How to organize an online conference. Nat Rev Mater 5:253–6 (2020).

27. Achakulvisut T, Ruangrong T, Bilgin I, Van Den Bossche S, Wyble B, Goodman DFM, et al. Point of View: Improving on legacy conferences by moving online. eLife 9, e57892 (2020). Available from: https://elifesciences.org/articles/57892

28. Royal Society of Chemistry Twitter Poster Conference 2019. Available from: http://www.rsc.org/events/detail/37540/rsc-twitter-poster-conference-2019

29. How to participate in a Twitter Conference? (2020). Available from: https://44c203da-d912-4bb2-9746-daaf050732b4.filesusr.com/ugd/11d9ac_9bae98d91a384fdfb6807233f1e9188f.pdf

30. Data Science Slack Communities to Join-Reach out in Slack to level up in your career. Available from: https://towardsdatascience.com/15-data-science-slack-communities-to-join-8fac301bd6ce

31. Brain Web: a permanent virtual space for online collaborations on projects related to neuroscience (2020). Available from: https://brain-web.github.io

32. Sever R, Roeder T, Hindle S, Sussman L, Black KJ, Argentine J, Manos W, Inglis JR. bioRxiv: the preprint server for biology. bioRxiv (2019); Available from: https://www.biorxiv.org/content/10.1101/833400v1

33. Hindle S, Saderi D. PREreview — a new resource for the collaborative review of preprints. eLife (2017); Available from: https://elifesciences.org/labs/57d6b284/prereview-a-new-resource-for-the-collaborative-review-of-preprints

34. Sanz-Cobena A, Alessandrini R, Bodirsky BL, Springmann M, Aguilera E, Amon B, et al. Research meetings must be more sustainable. Nat Food 1:187–189 (2020). Available from: https://www.nature.com/articles/s43016-020-0065-2?proof=trueIn%25EF%25BB%25BF

35. Tamar Snyder. Planning a Conference? Add a Lactation Room to Your Checklist (2014). Available from: https://ejewishphilanthropy.com/planning-a-conference-add-a-lactation-room-to-your-checklist/

36. Calisi RM and a Working Group of Mothers in Science. Opinion: How to tackle the childcare-conference conundrum. Proc Natl Acad Sci USA 115(12):2845–9 (2018). Available from: https://www.pnas.org/content/115/12/2845?platform=hootsuite

37. Foxx AJ, Barak RS, Lichtenberger TM, Richardson LK, Rodgers AJ, Williams EW. Evaluating the prevalence and quality of conference codes of conduct. Proc Natl Acad Sci USA 116(30):14931–6 (2019). Available from: https://www.pnas.org/content/116/30/14931

38. Favaro B, Oester S, Cigliano JA, Cornick LA, Hind EJ, Parsons ECM, Woodbury TJ. Your Science Conference Should Have a Code of Conduct. Front Mar Sci. 3(103):4 (2016).

39. Abbott A. Biologist found dead during Crete conference (2019).; Available from: https://www.nature.com/articles/d41586-019-02132-3

40. Chand D, Nayak S, Bhat KS, Parikh S, Singh Y, Kamath AA. A mobile application for Women’s Safety: WoSApp. IEEE (2016). TENCON, IEEE Region 10 International Conference. Available from: https://ieeexplore.ieee.org/document/7373171/authors#authors

41. Air Travel Mitigation Fund-University of California Los Angeles (2020). Available from: https://www.sustain.ucla.edu/wp-content/uploads/Air-Travel-Mitigation-Fund-Program-Guidelines-1.pdf

42. Reducing CO2 and air travel at ETH Zurich. ETH Zurich mobility platform. Available from: https://ethz.ch/content/dam/ethz/associates/services/organisation/Schulleitung/mobilitaetsplattform/ETH0041_Flugreisen_Factsheet_EN_f03.pdf

43. Code of Conduct to support a low-carbon research culture. Tyndall Travel Strategy - towards a culture of low carbon research for the 21st Century (2019). Available from: https://tyndall.ac.uk/travel-strategy

44. Evaluating Computer Scientists and Engineers For Promotion and Tenure-Computing Research Association. Available from: https://cra.org/resources/best-practice-memos/evaluating-computer-scientists-and-engineers-for-promotion-and-tenure/

45. Guidelines on the criteria for promotion and tenure-University of Minnesota College of Science and Engineering. Available from: https://cse.umn.edu/college/guidelines-criteria-promotion-and-tenure

46. University College London Travel Emissions (2020). Available from: https://www.ucl.ac.uk/sustainable/travel-emissions

47. Anderson K. The inconvenient truth of carbon offsets. Nature 484,7 (2012). Available from: https://www.nature.com/news/the-inconvenient-truth-of-carbon-offsets-1.10373

48. Popkin G. How much can forests fight climate change? Nature 565:280–2 (2019). Available from: https://www.nature.com/articles/d41586-019-00122-z

49. Huang J, Gates AJ, Sinatra R, Barabási A-L. Historical comparison of gender inequality in scientific careers across countries and disciplines. Proc Natl Acad Sci 117(9):4609– 16 (2020). Available from: http://www.pnas.org/lookup/doi/10.1073/pnas.1914221117

50. Shannon M Ruzycki SF, Madalene Earp AB, Kirstie C Lithgow. Trends in the Proportion of Female Speakers at Medical Conferences in the United States and in Canada, 2007 to 2017. JAMA Netw Open 2(4):e192103 (2019). Available from: https://jamanetwork.com/journals/jamanetworkopen/fullarticle/2730476

51. 2019 Women, Minorities, and Persons with Disabilities Report. National Science Foundation (2019). Available from: https://www.nsf.gov/news/news_summ.jsp?cntn_id=297944&org=NSF&from=news

52. Mervis J. New Answers for Increasing Minorities in Science. Science(2010); Available from: https://www.sciencemag.org/news/2010/09/new-answers-increasing-minorities-science

53. Inclusive Scientific Meetings-500 Women Scientists. Available from: https://500womenscientists.org/inclusive-scientific-meetings

54. Martin JL. Ten Simple Rules to Achieve Conference Speaker Gender Balance. PLOS Comput Biol (2014). Available from: https://journals.plos.org/ploscompbiol/article?id=10.1371/journal.pcbi.1003903

55. Resources for Reproducible Research. Available from: https://www.repro4everyone.org/

56. The Carbon Footprint of Academic Conferences: Evidence from the 14th EAAE Congress in Slovenia-Agricultural Economics Society and European Association of Agricultural Economists (EAAE). EuroChoices 15(2):56–61 (2015). Available from: https://onlinelibrary.wiley.com/doi/pdf/10.1111/1746-692X.12106

57. Aron AR, Ivry RB, Jeffrey KJ, Poldrack RA, Schmidt R, Summerfield C, et al. How Can Neuroscientists Respond to the Climate Emergency? Neuron 106(1):17–20 (2020). Available from: https://www.sciencedirect.com/science/article/pii/S0896627320301422

58. Mair J. Profiling conference delegates using attendance motivations. J Conv Event Tour 11(13):176–94 (2010). Available from: https://www.tandfonline.com/doi/abs/10.1080/15470148.2010.502032

59. Wynes S, Donner SD. Addressing Greenhouse Gas Emissions from Business-Related Air Travel at Public Institutions: A Case Study of the University of British Columbia. Department of Geography, University of British Columbia (2018). Available from https://pics.uvic.ca/sites/default/files/AirTravelWP_FINAL.pdf

60. Heike J. Transnational academic mobility and gender. Glob Soc Educ 9(2):183–209 (2011). Available from: https://www.tandfonline.com/doi/full/10.1080/14767724.2011.577199

61. 15th International Conference on Music Perception and Cognition (ICMPC15)-Graz Austria (2018). Available from: https://music-psychology-conference2018.uni-graz.at/en/about/

62. Freeman RB, Huang W. Collaborating with People Like Me: Ethnic Coauthorship within the United States. S289–318 (2013). Available from: https://www.journals.uchicago.edu/doi/abs/10.1086/678973

63. Gernsbacher MA. Video Captions Benefit Everyone. Policy Insights Behav Brain Sci. 2(1):195–202 (2015). Available from: https://journals.sagepub.com/doi/10.1177/2372732215602130

64. Wickham H. ggplot2: elegant graphics for data analysis (2016). Available from: https://books.google.com/books?hl=en&lr=&id=XgFkDAAAQBAJ&oi=fnd&pg=PR8&ots=spY07U8X3P&sig=Vw5aHFonM3Ee56OTEuWCfgUXA-c#v=onepage&q&f=false

65. Neuwirth E. RColorBrewer: ColorBrewer Palettes. The R Foundation (2014). Available from: https://cran.r-project.org/web/packages/RColorBrewer/index.html

66. Sarabipour S. Research Culture: Virtual conferences raise standards for accessibility and interactions. eLife 9, e62668 (2020). Available from: https://elifesciences.org/articles/62668

