## Supplementary File_Sarabipour et al_2021 for "Evaluating features of scientific conferences: A call for improvements"

### Supplementary Methods

#### An Online Database of in-person conferences in various scientific disciplines

**Data extraction: Generating the list of in-person conferences.** The majority of academic conferences do not readily report any form of raw or aggregate meeting statistics before or after the event. To compile a large list of national and international in-person conferences, we leveraged Google and lists available online (Table S2) and searched for conferences in a wide selection of fields and looked for conferences that scientists would be commonly attending. We also used a number of conference lists compiled by journals and learned societies available online. All the academic meetings in our database were held between January 2018 and December 2019 and spanned a wide range of disciplines in engineering, natural sciences, life and biomedical sciences, medicine and a number of humanities disciplines (Table S3). The data collected represented conferences organized by over 150 scientific societies and other organizations. We also selected conferences organized by funding agencies such as the United States National Institutes of Health (NIH), United Kingdom Wellcome Trust and United States National Science Foundation (NSF), conferences organized by publishing industry journals such as Nature and Elsevier as well as conferences organized by diagnostic and therapeutic biological companies such as pharmaceutical corporations. We recorded the name and geographic location of the conferences in our manually collated database available online on (https://elifeambassadors.github.io/improving-conferences/). Google searches were performed for each conference to identify the learned society/organization/entity/university/corporation in charge of organizing the meeting and the number of years the meeting has been held and the frequency of the meeting (i.e. held annually, biennially, more or less frequent). Further searches were performed to find the exact or estimated number of members each scientific society (if applicable to a specific conference) had registered. A separate search was performed for each conference, on the meeting website to identify the number of meeting attendees (exhibitors included). If the number of attendees was not provided on the meeting website, full program or program report (in the cases where the conference was already held), a Google search was performed instead. If after all search avenues were exhausted, the exact attendee number was not available online, an estimated number of attendees or society members was used instead (by comparison to members and attendees counts of societies in related fields). The number of attendees was collected to calculate/estimate the total meeting air travel and other (ground transportation, food and accommodation) attendee footprint. We quantified various features of the conferences from the information we had collected manually using the corresponding conference website. When quantifying the number of men and women speakers, chairs, organizing and scientific committee members, in cases where a name appeared ambiguous, we searched for the first and last name of the researcher with their university affiliation and field of expertise/research/conference topics to locate a photo. Aggregate summaries of the database are presented as Figure 1 and supplementary Tables S2-S11. On each conference website, the name of the conference, geographic location of the meeting, hosts/funders/sponsors, the year that the meeting was held, number of attendees, registration costs for different types of attendees, digital or virtual conferences with live streaming of talks and digital poster options (meaning online posters, excluding digital or ePosters on physical stands on site), availability of recorded talks and ePosters/iPosters (digital libraries), digital libraries/archives of talks, posters and slides after the meeting was held, total conference cost for each attendee (converted to US dollars), minimum and maximum meeting registration cost for each attendee availability of full (both oral and poster sessions detailed) conference program as electronic/virtual meeting scientific program, (keywords “program”, “agenda”, “schedule” book, “meeting planner”, “meeting App” for the scientific program book file or App (talks and abstracts), estimate air travel and other carbon footprint for total attendees of each meeting, on-site or off-site, free or at a cost childcare facilities, lactation or nursing rooms or personal consideration rooms, caregiver grants, types of career development workshops offered, types of Early career Researcher (ECR) promotion events (poster or oral/platform presentation awards, ECR symposia), networking events (mixers, meet and greet, ice-breaker or meet the experts events for trainees), availability of code of conduct, code of ethics (or research integrity), local safety instructions or Apps or facilities (e.g. a body system for watching out for fellow attendees, free bus/train), gender equity statement, diversity statement, invited speakers (those hand-picked by conference organizers, rather than scientists who put themselves forward to give talks or present posters) such as keynote speaker/opening/closing speakers names, plenary speaker names, conference chair names, organizing or steering committee names, session chair/organizer/moderator/convenor/discussion leader names and invited or featured speaker names, environmental sustainability events, public outreach events (such as free public lectures or engagement with local schools and libraries) and green policies were indicated and recorded in our database for this study. To explore early career promotion and appreciation events on conference websites, we looked with terms such as “award”, “scholarship”, “fellowship”, “funding” and “prize”. To search for session heads, we searched with keywords such as “session chair”, “session moderator”, “session organizer” and “session convenor”. To find the conference organizing committee, searches were performed with words such as “program committee” or “scientific committee”. Searches were also performed to count men and women members of the “organizing committee” or “planning committee” or “steering committee”. A separate Google search provided the corresponding academic discipline, the scientific organization associated or supporting the meeting and the most recent number of society members. To find information about sponsoring organizations, institutions or corporations which provided financial, intellectual or media support/partners, searches were performed using keywords “sponsors”, “supporters” and “partners” on each meeting website. To source information on code of ethics, we searched the conference and associated learned society webpage with the keywords “code of ethics”, “statement of ethics”, “professional code of conduct” and “statement/code of research integrity”. To explore the available information on the statement of gender equity, we searched the conference and corresponding society website with the keywords, “gender balance statement”, “gender equity statement”, “diversity statement”, “inclusion statement”. We also examined the conference webpages for their “sustainability initiatives” or “green policies strategies or considerations” incorporated into the organization of these conferences. We also included the meeting URL address and other important announcements, features or notices available for attendees (such as epidemics or phishing scams for flights and accommodations that could impact attendees).

### Data Visualization

Data visualized in Figure 2 were sourced from the Carbon Footprint Calculator: https://calculator.carbonfootprint.com/calculator.aspx?tab=3. Data visualized in Figure S3 on the annual fossil CO2 emission per capita for the year 2017 were sourced from https://ourworldindata.org/co2-and-other-greenhouse-gas-emissions and https://www.epa.gov/ghgemissions/global-greenhouse-gas-emissions-data. Data visualized in Figure S1 on the 2019 gross minimum wage worldwide (full amount an employer pays before taxes and other deductions are withheld) minimum annual (earned for 12 months in US dollars) were sourced from: https://data.worldbank.org/indicator/PA.NUS.PRVT.PP. Data visualized in Figure 3 and S2 were sourced from: https://www.henleyglobal.com/henley-passport-index/. All supplementary figures were made with R and the ggplot2 package (https://books.google.com/books?hl=en&lr=&id=XgFkDAAAQBAJ&oi=fnd&pg=PR8&ots=spY07U8X3P&sig=Vw5aHFonM3Ee56OTEuWCfgUXA-c#v=onepage&q&f=false), with colours from the RcolorBrewer package (https://cran.r-project.org/web/packages/RColorBrewer/index.html). Code and data underlying all figures is available on (http://doi.org/10.5281/zenodo.4067243). All in-person conference data displayed in Figure 1 were manually collated at https://elifeambassadors.github.io/improving-conferences/.

**Table S1. American Chemical Society Fall & Spring in-person Annual Meeting Attendee Statistics**

| Year | Location | Attendees | Students | Expo-Only | Exhibitors | Total |
|---|---|---|---|---|---|---|
| Spring 2019 | Orlando, FL | 7,974 | 6,043 | 444 | 858 | 15,754 |
| Fall 2018 | Boston, MA | 8,380 | 3,691 | 682 | 1,172 | 14,463 |
| Spring 2018 | New Orleans, LA | 9,067 | 6,472 | 334 | 879 | 16,752 |
| Fall 2017 | Washington, DC | 8,400 | 2,999 | 477 | 1,068 | 12,944 |
| Spring 2017 | San Francisco, CA | 9,830 | 6,920 | 967 | 1,200 | 18,917 |
| Fall 2016 | Philadelphia, PA | 7,860 | 3,257 | 697 | 1,175 | 12,989 |
| Spring 2016 | San Diego, CA | 8,776 | 5,989 | 477 | 1,068 | 16,310 |
| Fall 2015 | Boston, MA | 8,599 | 3,468 | 595 | 1,266 | 13,928 |
| Spring 2015 | Denver, CO | 7,612 | 5,142 | 357 | 847 | 13,958 |
| Fall 2014 | San Francisco, CA | 10,372 | 3,724 | 550 | 1,128 | 15,774 |
| Spring 2014 | Dallas, TX | 7,083 | 5,172 | 433 | 810 | 13,498 |

| | | | | | | |
|---|---|---|---|---|---|---|
| Fall 2013 | Indianapolis, IN | 6,849 | 2,664 | 418 | 872 | 10,803 |
| Spring 2013 | New Orleans, LA | 8,329 | 5,848 | 383 | 913 | 15,473 |
| Fall 2012 | Philadelphia, PA | 8,130 | 3,184 | 690 | 1,224 | 13,228 |
| Spring 2012 | San Diego, CA | 9,266 | 5,749 | 716 | 1027 | 16,758 |
| Fall 2011 | Denver, CO | 6,712 | 2,387 | 356 | 998 | 10,453 |
| Spring 2011 | Anaheim, CA | 7,641 | 4,688 | 596 | 1,097 | 14,022 |
| Fall 2010 | Boston, MA | 8,554 | 3,240 | 770 | 1,508 | 14,072 |
| Spring 2010 | San Francisco, CA | 10,200 | 5,717 | 927 | 1,223 | 18,067 |
| Fall 2009 | Washington, DC | 8,914 | 3,159 | 671 | 1,449 | 14,193 |
| Spring 2009 | Salt Lake City, UT | 5,963 | 3,443 | 420 | 780 | 10,606 |
| Fall 2008 | Philadelphia, PA | 8,699 | 2,972 | 838 | 1,488 | 13,997 |
| Spring 2008 | New Orleans, LA | 7,581 | 4,671 | 385 | 1,158 | 13,795 |
| All years Combined | Total Attendees | 190,791 | 100,599 | 13,183 | 25,208 | 330,754 |
| Total Carbon Footprint | Average $CO_2$ produced:<br><br>Average Arctic ice melted: | ~190,791metric tons<br><br>572,373 m$^2$ | ~100,599 metric tons<br><br>301,797 m$^2$ | ~13,183 metric tons<br><br>39,414 m$^2$ | ~25,208 metric tons<br><br>75,624 m$^2$ | ~300,754 metric tons<br><br>902,262 m$^2$ ~ Over 4 times the area of NY Grand Central Station |

**Table S1.** Over a decade of American Chemical Society (ACS) annual meeting attendance has left a large carbon footprint. These are statistics for fall and spring annual meetings only and do not include the many in-person regional meetings of this society at various states across the United States (Data source (American Chemical Society National Meeting and Expo-Attendee Demographics (2019): https://www.acs.org/content/acs/en/meetings/national-meeting/exhibitors/attendee-demographics.htm). A round-trip flight from New York to San Francisco emits about 1 metric tons of carbon dioxide per person. Every metric ton of $CO_2$ emitted leads to 3 square meters of Arctic sea ice loss. If every attendee trip, food and accommodation produced about 1 metric tons of $CO_2$ on average, total ACS annual meeting attendance resulted in loss of over 900,000 m$^2$ of arctic ice. The Grand Central Station in New York, NY, the largest train station in the world by platform count, measures 200,000 square meters (m$^2$) in total area.

| **Table S2. Online sources with compiled lists of conferences** | | |
|---|---|---|
| **Type** | **Name** | **Website** |
| Conference List | Nature Ecology and Evolution | www.nature.com/natecolevol/about/conferences |

| | | |
|---|---|---|
| Conference List | the National Invasive Species Information Center | www.invasivespeciesinfo.gov/conferences-and-events |
| Journal List | List of ecology, evolution, and conservation journals | https://docs.google.com/spreadsheets/u/1/d/1uG2Dg0LogysCSAsK51Rh9lD_dxRRFOeo2jq92_TwqF0/htmlview |
| Broad Search platform | Google | www.Google.com |

**Table S2.** Lists and resources used to compile our conference database (4).

| **Table S3. Conference Disciplines** | |
|---|---|
| Physics, Astronomy, Space Research, Astronautics, Condensed Matter Physics | (3/270) |
| Chemistry, Chemical Engineering, Mass Spectrometry, Glycobiology | (7/270) |
| Molecular Biology, Molecular Life Sciences, Biochemistry | (4/270) |
| Microbiology, Microbial Communication, Microbial Population Biology, Veterinary Microbiology | (12/270) |
| Cell Biology, Single Cell Biology, Developmental Cell Biology, Mechanobiology | (10/270) |
| Developmental Biology, Zebrafish Biology, Evolutionary Developmental Biology | (3/270) |
| Behavioral Biology, Behavioral Ecology | (2/270) |
| Fish & Wildlife science, Forest insect science, Forest conservation | (3/270) |
| Coral Reef Biology, Island Biology | (2/270) |
| Land Ecology, Ecosystem Restoration, Lake Ecology, Soil & Water Conservation | (6/270) |
| Agriculture, Agricultural Engineering, Enology | (2/270) |
| Integrative & Comparative Biology | (1/270) |
| Veterinary Microbiology, Veterinary parasitology | (2/270) |
| Evolution, Human Evolution, Ecology, Archaea Ecology, Molecular Ecology, Microbial Ecology, Mathematics of Ecology, Ecology & Cancer | (44/270) |
| Zoology, Ornithology, Primatology, Mammalogy, Herpetology | (8/270) |
| Insectology, Entomology, Myrmecology, Pesticide Resistance, Beekeeping | (7/270) |
| Palaeontology, Palaeoanthropology, Vertebrate Palaeontology | (4/270) |
| Oceanography, Marine Biology, Ocean Science, Marine Mammalogy | (4/270) |
| Climate Science | (2/270) |

| | |
|---|---|
| Human Genetics, Fungal Genetics | (7/270) |
| Vertebrate Pest Science | (1/270) |
| Invertebrate Pathology | (1/270) |
| Epigenetics | (2/270) |
| Pathology, Molecular Pathology | (2/270) |
| Pharmacology & Drug Development, Neuropsychopharmacology, Pharmaceutical Sciences, Peptide Science, Redox Chemistry & Medicine | (5/270) |
| Mathematics, Mathematical & Theoretical Biology | (2/270) |
| Immunology, Virology, Infectious Disease, Clinical Microbiology, HIV, Vaccines, Allergy, Leukocyte Biology, Lymphology | (19/270) |
| Biophysics, X-ray Diffraction Crystallography & Scattering, Chromatin Biophysics | (5/270) |
| Neuroscience, Neurological Disorders | (3/270) |
| Cognitive Sciences | (3/270) |
| Plant Science (Plant-microbe interactions, bacterial wilt, aquatic plant science)/Synthetic Biology, Phytobiome research | (14/270) |
| Biomedical Engineering, Medical Physics, Biomedical Electronic Devices | (3/270) |
| Imaging, Microscopy | (5/270) |
| Biomolecular Research Facilities | (1/270) |
| Materials Science & Engineering, Computational Modelling of Materials, Mining Engineering | (5/270) |
| Mechanotronic & Robotics | (1/270) |
| Environmental & Energy Engineering | (1/270) |
| Systems Biology, Computational Biology, Molecular Dynamics | (11/270) |
| Bioinformatics, Data science, Machine Learning | (2/270) |
| Computer Science (Software & Hardware Applications) | (3/270) |
| Conservation Biology | (5/270) |
| Cancer Research | (1/270) |
| Epidemiology, Molecular Epidemiology | (2/270) |
| Bone & Skeletal Disorders, Autism, Osteoporosis, Osteoarthritis, Musculoskeletal Disease | (4/270) |
| Psychology, Psychonomics | (2/270) |
| Clinical Laboratory Science, Clinical Research | (2/270) |
| Nutrition | (1/270) |

| | |
|---|---|
| Toxicology | (1/270) |
| Ophthalmology | (1/270) |
| Geology & Earth Science, Biogeography (invasive species), Geography | (6/270) |
| Health care/Human health, Medicine, Anaesthesiology | (3/270) |
| Cardiovascular Disease, Diabetes, Vascular Biology | (3/270) |
| Alzheimer's | (1/270) |
| Endocrinology | (1/270) |
| Maternal Medicine, Infant studies (Development of infants), Reproductive Biology, Study of birth defects, Pediatric Research | (5/270) |
| Rare Disease | (1/270) |
| Gerontology, Biology of Aging | (1/270) |
| Sports Medicine | (1/270) |
| Behavioral Medicine | (1/270) |
| Linguistics | (1/270) |
| History, Philosophy & Social Sciences of Biology, Ethnobiology, History of Medicine | (4/270) |
| Biosafety & Biosecurity | (1/270) |
| Education, Gender, Diversity (Research Culture) | (2/270) |

**Table S3.** The research disciplines for the 270 conferences analyzed. Conference names and year the meeting was held are available in the database (4).

| **Table S4. Frequency of in-person National & International Conferences Worldwide** | |
|---|---|
| Annual | 73.7% (199/270) |
| Biennial | 20% (54/270) |
| Triennial | 1.5% (4/270) |
| Quadrennial | 3.7% (10/270) |
| Other: meetings held multiple times a year | 1.1% (3/270) |

**Table S4.** The frequency of occurrence of the 270 conferences examined in this study (4).

| Table S5. Number of years in-person conferences have been held | |
|---|---|
| Minimum | 1 Year |
| Average | 29 Years |
| Maximum | 187 Years |

**Table S5.** The number of years the 270 academic conferences examined were held (4).

| Table S6. Number of Conference Attendees (Researchers + Exhibitors | |
|---|---|
| Minimum | 70 researchers |
| Average | 2,500 researchers |
| Maximum | 31,000 researchers |
| Estimate total Attendees | ~859,114 researchers |
| Estimate total Learned Society Members, representing over 150 learned societies | ~1,658,602 researchers |

**Table S6.** The number of attendees varied widely for the 270 conferences in our database (4).

| Table S7. Cost of Conference Registration for Attendees | |
|---|---|
| Minimum | Free of charge |
| Average | Over US$200 |
| Maximum | US$2,296 |
| Average Total Funds spent for all attendees (~859,114 researchers, each on average spent US$1500) | US$1.288 billion |

**Table S7.** The registration cost for the 270 conferences we examined. Minimum and maximum registration costs were recorded for each conference, available online (4).

| Table S8. Travel Carbon Footprint of Conference Attendees | |
|---|---|
| Minimum per attendee | Traveling 1 mile by train to the convention center produces 0.5 kg $CO_2$ |
| Maximum per attendee | Flying from Perth, Australia to London, United Kingdom and back for Virology & Infectious Diseases (ICVID) 2019 generated about 3,153 kg (3.47 tons) of $CO_2$ |
| Total **Air Travel** carbon footprint generated from 270 conferences (~859,114 attendees combined, average air travel $CO_2$ of 2 tons per attendee) | 1,718,228 tons of $CO_2$ |
| Total **Other** carbon footprint generated from 270 conferences (~859,114 attendees combined, average all other $CO_2$ production of 0.5 tons per attendee) | 429,557 tons of $CO_2$ |
| Aggregate Attendee travel carbon footprint for a trip to the American Association of Geographers annual meeting | Air travel by 6,741 attendees to and from a single meeting of the American Association of Geographers in Seattle produced ~16,000 metric tons of $CO_2$, equivalent to the amount that 53,500 people living in Haiti generated during 2014: https://www.tandfonline.com/doi/full/10.1080/00330124.2013.784954 |
| Further examples & resources on air travel and researcher carbon footprint | Observed Arctic sea-ice loss directly follows anthropogenic $CO_2$ emission (2016): https://science.sciencemag.org/content/354/6313/747<br><br>American Physical Society meeting statistics (2012): https://www.aps.org/publications/apsnews/201204/lrgestmarchmeet.cfm<br><br>atomsfair-Climate-friendly air travel: https://www.atmosfair.de/en/offset/flight/<br><br>Carbon Footprint Calculator: https://calculator.carbonfootprint.com/calculator.aspx?tab=3<br><br>CO2 emissions (metric tons per capita)-The World Bank: https://data.worldbank.org/indicator/EN.ATM.CO2E.PC<br><br>An IPCC special report on the impacts of global warming of 1.5 °C above pre-industrial levels and related global greenhouse gas emission pathways. New York, NY: The Intergovernmental Panel of United Nations on Climate Change (2018): https://www.ipcc.ch/sr15/ |

| | |
|---|---|
| | Reducing emissions from aviation. European Union: European Commission Climate Action (2019): https://ec.europa.eu/clima/policies/transport/aviation_en<br><br>The climate mitigation gap: education and government recommendations miss the most effective individual actions (2017): https://iopscience.iop.org/article/10.1088/1748-9326/aa7541<br><br>Carbon footprint of science: More than flying (2013): https://www.sciencedirect.com/science/article/pii/S1470160X13002306<br><br>How your flight emits as much $CO_2$ as many people do in a year (2019): https://www.theguardian.com/environment/ng-interactive/2019/jul/19/carbon-calculator-how-taking-one-flight-emits-as-much-as-many-people-do-in-a-year<br><br>Early Human Health Effects of Climate Change-WHO Office for Europe (1998): http://www.euro.who.int/__data/assets/pdf_file/0006/119184/E64599.pdf<br><br>Transformative change requires resisting a new normal (2020): https://www.nature.com/articles/s41558-020-0712-5<br><br>Report Of The Secretary-General On The 2019 Climate Action Summit-The Way Forward In 2020: https://www.un.org/en/climatechange/assets/pdf/cas_report_11_dec.pdf<br><br>Code of Conduct to support a low-carbon research culture: Tyndall Travel Strategy - towards a culture of low carbon research for the 21st Century (2019): https://tyndall.ac.uk/travel-strategy<br><br>Addressing Greenhouse Gas Emissions from Business-Related Air Travel at Public Institutions: A Case Study of the University of British Columbia. Department of Geography, University of British Columbia (2018): https://pics.uvic.ca/sites/default/files/AirTravelWP_FINAL.pdf |

**Table S8.** Examples of individual $CO_2$ production and aggregate amount for 270 conferences held between 2018 and 2019. Details of $CO_2$ production for each meeting is detailed in the online database (4).

| **Table S9. Summary key considerations of the 270 Scientific Conferences Analyzed** | | | |
|---|---|---|---|
| **Key Considerations** | | **Yes** | **No** |
| Equity, Intersectionality & Inclusivity | Live streaming or Recordings of **some or all** talks is available at the time of in-person meeting | 3.7% (10/270) | 96.3% (260/270) |

| | | | |
|---|---|---|---|
| Considerations | Live streaming or Recordings of **ALL** talks is available at the time of in-person meeting | 3.3% (9/270) | 96.7% (261/270) |
| | Virtual Reality or Digital Posters (e-Posters/ i-Posters/Twitter Posters) | 1.4% (4/270) | 98.6% (266/270) |
| | Archives of all or most recorded talks or e-posters available from conference or scientific society website from previous meeting years | 11% (30/270) | 89% (240/270) |
| | Code of Conduct (COC) | 41% (111/270) | 59% (159/270) |
| | Code of Research Ethics & Integrity | 22% (59/270) | 78% (211/270) |
| | On-site Facilities for Mothers such as Lactation/Nursing Room also called a Personal Considerations Room | 15% (41/270) | 85% (229/270) |
| | Caregiver Grants | 12.2% (34/270) | 87.8% (236/270) |
| | Some form of Childcare (free or at cost) available on-site | 19% (51/270) | 81% (219/270) |
| | Free on-site Childcare | 11% (30/270) | 89% (240/270) |
| | On-site Childcare at a cost ($$) | 8% (21/270) | 92% (249/270) |
| | ECR Training Workshops for Career Development | 35% (94/270) | 65% (176/270) |
| | ECR promotion Events (special symposia, talks, poster sessions for postdoctoral researchers on the job market, ECR awards) | 38.5% (104/270) | 61.5% (166/270) |
| | ECR Networking Events (such as mixer/ice-breaker events with fellow ECRs and senior researchers) | 20% (54/270) | 80% (216/270) |
| | ECR travel Award (Limited in number: 5-20 awarded during each conference) | 55% (149/270) | 45% (121/270) |
| | Local Safety Apps or Instructions for attendee physical safety in town/city | 4% (12/270) | 96% (258/270) |
| | Diversity statement<br>Reported on the conference website online | 22% (59/270) | 78% (211/270) |
| | Gender equity/balance statement<br>Reported on the conference website online | 8% (22/270) | 92% (248/270) |
| | Conferences chair gender balance<br>39.3% (106/270) of conferences reported this information on the meeting website | 46.2% (49/106) | 53.8% (57/106) |

| | | | |
|---|---|---|---|
| | Keynote speaker gender balance<br>41% (111/270) reported this information on the meeting website<br><br>Plenary speaker gender balance<br>47.4% (128/270) of conferences reported this information on the meeting website | 40.5%<br>(45/111)<br><br>38.3%<br>(49/128) | 59.5%<br>(66/111)<br><br>61.7%<br>(79/128) |
| | Invited/Featured speaker gender balance<br>52.2% (141/270) of conferences reported this information on the meeting website | 19.1%<br>(27/141) | 80.9%<br>(114/141) |
| | Session Chair gender balance<br>52.6% (142/270) of conferences reported this information on the meeting website | 36%<br>(51/142) | 64% (91/142) |
| | Organizing/Steering Committee gender balance<br>46.3% (125/270) of conferences reported this information on the reported website<br><br>Program/Scientific Committee gender balance<br>30% (81/270) of conferences reported this information on the reported website | 36.8%<br>(46/125)<br><br>27.2%<br>(22/81) | 63.2%<br>(79/125)<br><br>72.8% (59/81) |
| | Local public outreach events (e.g. public talks) | 6%<br>(15/270) | 94%<br>(255/270) |
| Environmental Sustainability Considerations | Sustainability Policy or Green Strategy<br>(e.g. buying carbon off-sets, going paperless, reducing plastic bottles, sourcing local vegetarian food options for catering) | 5.6%<br>(15/270) | 94.4%<br>(255/270) |
| | Electronic Apps or online program books (complete schedule of talks and posters/abstract book)<br>(in the form of interactive schedule or a .pdf file or mobile phone App) | 91%<br>(247/270) | 9% (23/270) |
| | Nature (e.g. Forest, Beach) clean-up walks/ events | 0% (0/270) | 100%<br>(270/270) |

**Table S9.** Summary of aggregate data from 270 conferences. Not every conference reported or designated scientists in certain roles. For instance, a number of conferences did not assign session chairs during their meeting. Plenary, keynote and invited/featured speaker roles were not also all consistently assigned in all meetings. Detailed data available on our conference online (4).

| **Table S10. Types of Educational/Career Development Workshops/Events Held** | | |
|---|---|---|
| **Type of Workshop** | **Number of Conferences which offered the workshop** | **Notes/Details** |

| Events/Workshops for Undergraduate/Graduate/Postdoctoral Trainees | | |
|---|---|---|
| Mentoring & Leadership Training | 23 | Overcoming bias through mentorship, mentoring connections, one-on-one Mentoring, Career Mentoring |
| Improving academic environments & culture | 1 | Research culture |
| Managing Yourself as a researcher | 1 | Leadership |
| Tools for Negotiations | 4 | Advocating for yourself and your brand/goals/<br>Advocating for your research publications |
| Professionalism: Building Success in Science | 1 | |
| Early career Researcher Networking & Conversations | 14 | Supporting ECRs/<br>Early career days/<br>ECR Challenges/<br>Community & Connections |
| Creating Early Career Researcher (ECR) committees | 1 | Engagement in scientific community |
| Effective Management of shared facilities | 1 | Management skills |
| Effective Teaching in Classrooms | 7 | Through practice/<br>Inclusive pedagogy/<br>Creating a comfortable and welcoming learning community: from a strategic syllabus to realized student engagement/ effective use of online resources for your class/<br>Developing a concept inventory to evaluate Student learning In undergraduate courses |
| Career Advice: Navigating the Job market | 11 | Career Transition/Hiring & Promotion/job skills/career planning for success/Diverse career paths in specific fields/First year on the jobs tips/career planning, getting hired, Searching, Applying, Interviewing, and Negotiating for Your First Job |
| Career Fair/Career Panel/Career Consultation | 10 | Building a good CV/<br>Recruitment event |
| Graduate & Postdoctoral researcher career development session | 12 | Networking: How to Create Your Dream Career, Networking, Informational Interviews for ECRs, navigating the path to |

| | | |
|---|---|---|
| | | professional success, preparing trainees for modern careers, IDPs: Individual Career Development plans |
| Developing a value statement | 4 | Elevator pitch, Three-minute thesis competition, Student & Early career pop talks (5-minute TED style talks) |
| Graduate & Postdoctoral Career Development | 8 | Developing graduate skills / The Strategic Postdoc: How to find & leverage your postdoc experience, creating an individual development plan/postdoctoral challenges/non-traditional postdocs, ECR discussions on various PhD projects with senior researchers, tips for successful graduate school applications, Career Speed-Networking Luncheon |
| Networking in research | 2 | |
| Engaging more Graduate & Postdoctoral trainees in scientific societies and their organizing committees | 3 | |
| Academic Career Track | 12 | Preparing your Written Application Materials: CV, Cover Letter, Research Statement, the job talk, Understanding the search process from the perspective of search committees and Decoding Job Announcements |
| Non-Academic Career Tracks | 17 | The industry interview, government & NGO jobs, Resume review session, progressive lunch with Industry |
| Writing Diversity Statements | 1 | |
| Scientific Communication | 23 | Increasing the impact of your research through social media, communicating for Impact: Workshop on engaging meaningfully with your neighbours, your elected officials, funders and the broader public about science/ turn your science into news/ effective communication of your data, How to design and give dynamic powerpoint talks and TED talks, scientific storytelling, Developing Strategies for |

| | | |
|---|---|---|
| | | Effective and Trustworthy Communication |
| Scientific Writing for Public | 5 | Uncomfortable conversations: Engaging Diverse Communities |
| Scientific writing in Academia | 10 | For trainees & New Principal Investigators, writing your paper, increasing the impact of your research, Writing from Qualitative Data |
| Public Policy for Scientists | 6 | How to transition your research to public policy, Advocating for Biomedical Research: we have done it so can you |
| Public outreach, Advocacy | 4 | Increasing your success and social capital in conservation |
| Citizen science workshops | 3 | Working with data collected by citizen scientists – challenges & opportunities, empowering citizen science leaders with tools for robust community engagement |
| Reproducibility in Research | 6 | Experimental or Computational research, Tools for open science: reproducible data analysis and paper writing in R |
| Data/Statistical Analysis Techniques | 2 | |
| Data Management | 8 | Data collection/<br>Database/<br>Repository Building Techniques/<br>Digital Tool development/How to use Github, Integrating Advanced Technologies to Improve Data Quality and Reduce Bias in Population Research and Management, Data Analytics, Data mining, Data Cleaning |
| Teaching computing tools to researchers | 3 | |
| Building Digital Data Collection Apps | 2 | Software Carpentry/<br>building Apps |
| Visualization or illustration Techniques | 6 | Graphic recording/<br>how to make a video of your research/artwork for your research publication, Data visualization |
| Grant proposal Writing | 17 | Optimizing Grant Applications/<br>Funding Opportunities, Training & Fellowship grants |

| | | |
|---|---|---|
| Grant information session with specific funding agencies, Funding & budgeting the research | 9 | e.g. NSF, NIH, FDA, European Council Funding Agency/ Early career grant writing opportunities, Communicating with Program Officers, study section review, progress reports, Science, Dollars, and Outcomes: The Critical Pieces of Budgeting You Can't Work Without, NIH Support for Typical and Non-typical Career Trajectories: getting to where you want to be |
| Forming Successful Collaborations, Networking | 7 | Core competencies for partnering)/ Navigating Team-science/ strategies to generate data/ joining consortium projects/ Collaborative Science with diverse stakeholders/ Industry collaborations & technology development/new technologies & expanded opportunities for collaboration, promoting yourself by making connections that count, finding your voice |
| Succeeding in Interdisciplinary Research, Effective Team-building and Communication workshop | 3 | |
| Multidisciplinary Research Needs | 1 | |
| International Opportunities in Science | 1 | Working as a scientist outside of the U.S. requires curiosity, adaptability, and open-mindedness, which are valuable qualities important for success in any career |
| Getting your foot at the door for leadership positions | 1 | Getting hired |
| Media Engagement/Communication | 3 | Media training |
| Public Involvement | 4 | Pitch your science to non-scientists/Public Engagement/ Citizen Science Data |
| Tips for STEM Educational Engagement outside Colleges & Universities | 1 | Engaging with schools, youth groups and home educators |
| Science & Arts | 2 | Researcher Film Festivals/Science & Poetry |
| Promoting Diversity, Equity & Inclusivity | 17 | In classrooms & In research environments/Promoting |

| | | |
|---|---|---|
| | | Women/LQBTQI researchers/cultural diversity, Diversity and Inclusion: Leveraging Actions Through Collaboration |
| Immigration Challenges | 1 | On Obtaining Visas |
| Mental Health & Well-Being | 1 | |
| Work-Life Balance | 4 | Avoiding burn out during your career |
| Responding to Bullying | 2 | |
| Gender inequalities in research environments | 3 | (e.g. in field work), Navigating power dynamics in academia |
| Implicit Bias, Bias Awareness in Academia | 3 | |
| Disability in Academia | 1 | |
| Imposter Syndrome | 1 | |
| Tips for successful publishing in a Journal | 11 | Publishing Q & A: Managing the Expectations of Peer Reviewers |
| Open Access (OA) publishing: Preprints/Peer Community In | 1 | (https://peercommunityin.org/) |
| Peer-reviewing Training | 8 | Reviewing training for reviewing manuscripts & grant applications, best practices, mock review sessions |
| Publication/Research Ethics/Teaching Integrity | 6 | Ethics in research environments, Professional Ethics & Advocacy, Irresponsible & wrong conduct of research |
| Fostering/Unlocking Early Career Potentials | 3 | Youth Capacity Building workshop |
| **Career Development Events/Workshops for Faculty** | | |
| Faculty Career Development Workshops<br>(PUI faculty) | 2 | |
| Early to Mid-Career PI challenges | 4 | New Faculty Forum/Striving for Success/Network, learn and find support, Tips for New and Early Stage Investigators:Planning for Success: Navigating Your First Faculty Position/How to get Tenure |
| PUI faculty training | 1 | Strategies for successful |

| | | |
|---|---|---|
| | | faculty/undergraduate student collaborative research at PUIs |
| Enabling Work–Life Balance in Your Research Group | 1 | |
| Meet the Editors | 9 | What editors expect in your publication/how to be a good associate editor |
| Lab Management Course for Principal Investigators (PIs) | 2 | including budget management, hiring staff, mentoring trainees |
| Grant Information Session for faculty and other professionals (Presented by Representatives of the funding agencies) | 2 | United States National Science Foundation information session |

**Table S10.** Details of the early- to mid-career trainee and PI development workshops offered only by 91 of the 270 conferences in our database (4). A number of conferences offered more than one career development workshop, workshops are categorized by topic.

| **Table S11. Types of Early Career Researcher (ECR) promotion events** | | |
|---|---|---|
| **Type of Event** | **Number of Conferences which offered the event** | **Notes** |
| **Promotion for Undergraduate/Graduate/Postdoctoral Trainees** | | |
| Graduate & Postdoctoral Trainee Reception/Mixer | 4 | Student Networking Events |
| Early Career trainee Research/Young Investigator Career Awards/Distinguished student Award/Merit Awards | 54 | For outstanding science, for ECRs with disabilities, for best talk, best poster, for women, minorities, best in their specific field (Graduate & Postdoctoral researchers), Infancy Early Career Researcher Award, Medical student achievement award |
| Trainee Career Development Award | 2 | |
| Technologists Award | 1 | Lab Technicians/non-doctoral research staff |
| Best talk/presentation/platform award | 22 | |
| Early career trainee symposium | 13 | Typically mini conference last 1-2 days |
| Networking | 12 | Meeting senior researchers at |

| | | |
|---|---|---|
| (meet & greet/ice-breaker/Mixer) events for ECRs | | conference breakfast, lunch or dinner events/student-faculty networking lunch, for under-represented minorities, LGBTQI trainees, for women, meet the women leaders, Student-Industry Mixer |
| Social reception following the Early Professionals Mini-Talk Symposium | 1 | Provides a chance to meet with the participants and other early career and mid/senior |
| First time delegate mentoring by other delegates | 1 | |
| Young Investigator Forum | 1 | |
| Outstanding Abstract Award | 1 | |
| Poster competition award | 24 | Meet the faculty candidate poster session/(best poster) or special viewing sessions |
| ECR job application networking event | 2 | Recruitment event |
| Fellow in training award | 3 | Basic research fellows, clinical research fellows, trainee award for innovation in medical education |
| Best paper published award | 4 | |
| Outstanding Dissertation/Thesis Award | 3 | |
| Participants favoriot talk or poster | 1 | |
| Diversity & Inclusion Awards | 3 | |
| Mobility Awards | 1 | |
| Science Communication Scholar Award | 1 | |
| Outreach initiative awards | 1 | |
| Ethics in research Essay and Video competition | 1 | |
| Best Essay Competition | 1 | |
| Student Engineering Design competition | 1 | |
| Undergraduate trainee platform presentations | 1 | |
| Undergraduate trainee | 6 | |

| | | |
|---|---|---|
| research awards | | |
| **Promotion efforts for ECR Faculty** | | |
| Early Career Research Awards for Principal Investigators (PIs) | 24 | Prizes for independent investigators/Faculty Development Award/ECR professionals for best research, R1 and PUI (Primarily Undergraduate Institutions) faculty, US-based or International are all eligible, Emerging leader award, Public policy award, Distinguished Early Career Contribution Award, ECR investigator Lectureship, basic researchers, Physician-scientists |
| Early Career Development Awards for Principal Investigators (PIs) | 1 | |
| Excellence in Research, Teaching & Service Award | 1 | |
| Mid-Career Research Awards | 1 | |
| Mid-Career Development Awards | 1 | |
| Emerging Leaders Mentorship Award | 2 | Mentoring Excellence Award |

**Table S11.** Details of the Early Career Researcher (ECR) promotion and networking events offered by a number of conferences in our database (4).

| **Table S12. List of additional examples & resources** | |
|---|---|
| **Theme** | **Additional Notes** |
| Human & financial resources devoted to R&D Worldwide | Facts and figures: human resources, the UNESCO Science Report, Towards 2030 (2019): https://en.unesco.org/node/252277<br><br>Appendix B. Determination of the minimum salary figure. In: The Postdoctoral Experience Revisited. Washington (DC): National Academies Press (US) (2014). (Committee to Review the State of Postdoctoral Experience in Scientists and Engineers; Committee on Science, Engineering, and Public Policy; Policy and Global Affairs; National Academy of Sciences; National Academy of Engineering; Institute of Medicine).<br><br>Average Postdoctoral Research Associate Salary in the United Kingdom. PayScale (2020): https://www.payscale.com/research/UK/Job=Postdoctoral_Research_Associate/Salary |

| | |
|---|---|
| ECR travel Grants | Travel Grants for Early Career Researchers at ECR Central: https://asntech.github.io/postdoc-funding-schemes/travel-grants/<br><br>Travel Grants for Latin American Early Career Researchers: https://ecrlarc.github.io/ |
| ECR Visas challenges | Visas for global health events—too many are losing their seat at the table: https://blogs.bmj.com/bmj/2019/07/30/ulrick-sidney-visas-for-global-health-events-too-many-are-losing-their-seat-at-the-table/<br><br>Visa denied? Navigating the visa minefield for visiting academics (2012): https://www.theguardian.com/higher-education-network/blog/2012/jan/12/academic-visa-research-south-africa |
| Gender Imbalance at Conferences | Representation of women among invited speakers at medical specialty conferences (2019): https://www.liebertpub.com/doi/full/10.1089/jwh.2019.7723?url_ver=Z39.88-2003&rfr_id=ori%3Arid%3Acrossref.org&rfr_dat=cr_pub%3Dpubmed&<br><br>Representation of women in speaking roles at surgical conferences (2019): https://www.sciencedirect.com/science/article/pii/S0002961019305987<br><br>Trends in the Proportion of Female Speakers at Medical Conferences in the United States and in Canada, 2007 to 2017 (2019): https://jamanetwork.com/journals/jamanetworkopen/fullarticle/2730476<br><br>Not “Pulling up the Ladder”: Women Who Organize Conference Symposia Provide Greater Opportunities for Women to Speak at Conservation Conferences (2016): https://journals.plos.org/plosone/article?id=10.1371/journal.pone.0160015 |
| Outreach activities at conferences | Outreach evenings at Ecology and Behavior Conference (2019): https://eb2019.sciencesconf.org/resource/page/id/9 |

| **Table S13. Examples of best practices & technologies used to improve scientific conferences** | |
|---|---|
| **Example 1** | Multi-Hub conferencing model: The attendees’ experience has been positive, showing that the multiple-site format can serve as an alternative to the traditional one-site format of holding national and international conferences: https://www.sciencedirect.com/science/article/pii/S0736585311000773<br><br>Scientific electronic panels can be held via platforms such as Virtual Keynote Symposia (VKS), Zoom, business version of Skype, YouTube streaming, Crowdcast, Vimeo livestream (https://livestream.com/), OBS Open Broadcaster Software (https://obsproject.com/). OBS Studio enables organizers to record talks.<br><br>ON24 platform-Webcasting and virtual event and environment technologies: https://www.on24.com/ |

| | |
|---|---|
| | Conferencing platform: https://www.sococo.com/<br><br>Conference Recording: https://slideslive.com/<br><br>Screen recorder & Video editor: https://screencast-o-matic.com<br><br>Confex used by Ecological Society of America annual virtual meeting 2020: https://confex.com/<br><br>Jitsi Meet: https://jitsi.org/jitsi-meet/ has decent functionality and ease of use. The Jitsi Meet Server is also open source https://github.com/jitsi/jitsi-meet.<br><br>Tencent Conferencing: https://intl.cloud.tencent.com/product/tcc<br><br>Conference software: https://slideslive.com/neurips<br><br>ShowMe: https://www.showme.com/<br><br>Networking App for events and conferences: https://whova.com/<br><br>Virtual Whiteboard: https://edu.google.com/products/jamboard/?modal_active=none<br><br>Discord: https://discord.com/<br><br>Discourse: https://www.discourse.org/<br><br>Reddit AMA/similar with authors after presentations: https://www.reddit.com/r/IAmA/<br><br>Meet by Google, GoToWebinar and GoToMeeting, or a combination of two of these platforms allowing both video conferencing and the opportunity to break into “breakout rooms” accommodating smaller discussion groups and one-on-one interactions (e.g. via Slack, Discord or Discourse). These platforms and others are being utilized for scientific chats on a daily and weekly basis where ideas are exchanged and collaborations are shaped among researchers:<br><br>15 Data Science Slack communities to Join-Reach out in Slack to level up in your career: https://towardsdatascience.com/15-data-science-slack-communities-to-join-8fac301bd6ce)<br><br>IEEE association offers tools such as WebEx (with recording option), Google meet, Google Hangouts (no recording option), Microsoft Teams, INXPO, YouTube for online meetings and recordings and tools such as Camtasia, QuickTime and WebEx Recorder for offline recordings.<br><br>For anonymous Q&A organizers can use platforms such as Slido: https://www.sli.do/ and Pigeonhole Live https://pigeonholelive.com/<br><br>Neuronmatch 2020 virtual conference: https://elifesciences.org/articles/57892 & https://neuromatch.io/ |
| **Example 2** | For example, traveling 40 miles to a regional conference results in emission of 20 kg of $CO_2$. Thus, where feasible, public or shared transportation options such as the bus or train should be used instead of a car or plane. |

| | |
|---|---|
| | American Physical Society (APS) regional meetings & American Chemical Society (ACS) organize multiple regional meetings throughout the year.<br><br>Evidence from the 14th Congress in Slovenia-Agricultural Economics Society and European Association of Agricultural Economists (EAAE): https://onlinelibrary.wiley.com/doi/pdf/10.1111/1746-692X.12106 ). |
| **Example 3** | Greenhouse gas emissions caused by business trips can make up 60% of the total university greenhouse gas emissions with air travel responsible for up to 94% of the institutional travel-related emissions.<br><br>Observed Arctic sea-ice loss directly follows anthropogenic CO2 emission (2016): https://science.sciencemag.org/content/354/6313/747<br><br>Reducing emissions from aviation. European Union: European Commission Climate Action (2019): https://ec.europa.eu/clima/policies/transport/aviation_en<br><br>The climate mitigation gap: education and government recommendations miss the most effective individual actions: https://iopscience.iop.org/article/10.1088/1748-9326/aa7541<br><br>Carbon footprint of science: More than flying (2013): https://www.sciencedirect.com/science/article/pii/S1470160X13002306<br><br>How your flight emits as much $CO_2$ as many people do in a year (2019): https://www.theguardian.com/environment/ng-interactive/2019/jul/19/carbon-calculator-how-taking-one-flight-emits-as-much-as-many-people-do-in-a-year<br><br>Academic Jet-Setting in a Time of Climate Destabilization: Ecological Privilege and Professional Geographic Travel (2013): https://www.tandfonline.com/doi/full/10.1080/00330124.2013.784954m<br><br>$CO_2$ emissions (metric tons per capita)-The World Bank: https://data.worldbank.org/indicator/EN.ATM.CO2E.PC<br><br>Addressing Greenhouse Gas Emissions from Business-Related Air Travel at Public Institutions: A Case Study of the University of British Columbia. Department of Geography, University of British Columbia (2018): https://pics.uvic.ca/sites/default/files/AirTravelWP_FINAL.pdf<br><br>Early Human Health Effects of Climate Change-WHO Office for Europe (1998): http://www.euro.who.int/__data/assets/pdf_file/0006/119184/E64599.pdf<br><br>Transformative change requires resisting a new normal (2020): https://www.nature.com/articles/s41558-020-0712-5 |
| **Example 4** | The Society for the Study of Evolution meeting sustainability strategies (2020), Cleveland, Ohio: https://www.evolutionmeetings.org/sustainability.html<br><br>MGAConf2018 conference name badge printed on seed paper-Australian Museums and Galleries Association WA: https://twitter.com/MuseumsAustWA/status/1003609563591933952 |

| | |
|---|---|
| **Example 5** | The UK Wellcome Trust, the US National Institutes of Health (NIH) & National Science Foundation (NSF) support conference organization via designated grants which conference chairs apply for in order to fund the meeting.<br><br>NIH Support for Scientific Conferences (R13 and U13): https://grants.nih.gov/grants/funding/r13/index.htm<br><br>NSF supports Research Coordination Networks (RCN): https://www.nsf.gov/funding/pgm_summ.jsp?pims_id=11691 |
| **Example 6** | iBiology: a project creating open-access science videos on biology research and science-related topics) has provided virtual talks and lectures for a number of years: https://www.molbiolcell.org/doi/10.1091/mbc.e14-02-0756<br><br>iBiology Virtual Talks (2020): https://www.ibiology.org/<br><br>The North American Vascular Biology Organization (NAVBO) holds a number of yearlong webinars, online journal clubs and online mini-symposia (2020): https://www.navbo.org/events/online |
| **Example 7** | Low-carbon, virtual science conference in multi-hub virtual mode (2019): https://www.nature.com/articles/d41586-019-03899-1<br><br>First Carbon-Reduced Chronobiology Conference-European Biological Rhythms Society (EBRS) (2019): https://srbr.org/join-the-first-carbon-reduced-chronobiology-conference-on-november-18/<br><br>The inaugural Photonics Online Meet-up (POM), the first all-online conference for photonics researchers (2020): https://sites.usc.edu/pom/<br><br>2nd Palaeontological Virtual Conference (2020): http://palaeovc.uv.es/<br><br>Systems Biology: Global Regulation of Gene Expression: a Virtual Meeting by Cold Spring Harbor Laboratory meetings & courses (2020): https://meetings.cshl.edu/meetings.aspx?meet=SYSTEMS&year=20<br><br>Society for Neuroscience (SfN) Virtual Conferences: https://www.sfn.org/Meetings/Virtual-Conferences<br><br>Neuromatch: An online conference in Computational Neuroscience (2020): https://neuromatch.io/<br><br>Neuronal Circuits: a Virtual Meeting. Cold Spring Harbor Laboratory (2020): https://meetings.cshl.edu/meetings.aspx?meet=CIRCUITS&year=20<br><br>CNS2020 Virtual: Cognitive Neuroscience Society Annual Meeting (2020): https://www.cogneurosociety.org/annual-meeting/<br><br>The age of the Twitter conferences: https://science.sciencemag.org/content/352/6292/1404.2<br><br>R-Ladies Global-a world-wide organization to promote gender diversity in the R community 2020: https://rladies.org/<br><br>5th International Electronic Conference on Medicinal Chemistry Online (2019): https://www.mdpi.com/journal/molecules/events/10089 |

| | |
|---|---|
| | Molecular Biosystems Conference 2019(#MBIOSYS19)-Eukaryotic Gene Regulation and Functional Genomics, Puerto Varas, Los Lagos, Chile (2019): http://www.molbiosystems.com/<br><br>15th International Conference on Music Perception and Cognition (ICMPC15)-Graz Austria (2018): https://music-psychology-conference2018.uni-graz.at/en/about/<br><br>A nearly carbon-neutral conference model- white paper/practical guide (2018): https://hiltner.english.ucsb.edu/index.php/ncnc-guide/#about |
| **Example 8** | 1st Cyanobacteria Twitter Conference, 24 October 2018 Australian Rivers Institute: https://cyanocost.wordpress.com/2018/08/27/1st-cyanobacteria-twitter-conference-24-october-2018/<br><br>Brain Twitter Conference 2019 #brainTC: Neuroscience making an impact: https://brain.tc/<br><br>Twitter for scientists: https://www.nature.com/articles/s41568-019-0170-4 |
| **Example 9** | Air Travel Mitigation Fund, University of California Los Angeles (2020): https://www.sustain.ucla.edu/wp-content/uploads/Air-Travel-Mitigation-Fund-Program-Guidelines-1.pdf |
| **Example 10** | There are multiple online, open access databases listing women and LGBTQIA ECR trainees and faculty which can be utilized for selection of conference speakers:<br><br>500 Women Scientists-Request a Woman Scientist: https://500womenscientists.org/ 500 Queer Scientists-Request a Scientist: https://www.500queerscientists.com/<br>Women in BrainStim database: http://womeninbrainstim.com/search/ |
| **Example 11** | Air travel by an academic in the UK who on average attends only one international conference or meeting per year by plane, will produce $CO_2$ emissions footprint of about five tons. This is over ten times as much as the average UK person's carbon footprint from leisure flights, and nearly 20% more than the average UK citizen's total annual carbon footprint from travel and home energy combined.<br><br>Who emits most? Associations between socio-economic factors and UK households' home energy, transport, indirect and total CO2 emissions: https://www.sciencedirect.com/science/article/pii/S0921800913000980<br><br>Why I will be flying less (2019): http://www.russpoldrack.org/2019/06/why-i-will-be-flying-less.html |
| **Example 12** | The cost of a recent academic conference of 20,000 attendees in Mexico reached over US$190 million: https://www.springer.com/la/book/9783540372233). |
| **Example 13** | Designing a Virtual Neuroscience Conference: Organizers mimic in-person networking using a matching algorithm (2020): |

| | |
|---|---|
| | https://www.simonsfoundation.org/2020/04/03/designing-a-virtual-neuroscience-conference/<br><br>How to moderate a crowdcast (neuro)science meeting (2020): https://medium.com/@kording/how-to-moderate-a-crowdcast-neuro-science-meeting-858afc8dfa05 |
| **Example 14** | Universal access to scientific and medical research via funder preprint mandates: https://journals.plos.org/plosbiology/article?id=10.1371/journal.pbio.3000273 |
| **Example 15** | American Chemical Society National Meeting & Expo-Attendee Demographics (2019): https://www.acs.org/content/acs/en/meetings/national-meeting/exhibitors/attendee-demographics.html |
| **Example 16** | The European Powder Diffraction Conference (EPDIC17) statement on Gender Balance, Sibenik, Croatia (2020): https://www.epdic17.org/registration<br><br>Gender Equity and Diversity, Australian Biophysical Society (2020): https://www.biophysics.org.au/meetings.html |
| **Example 17** | Code of Conduct for R Conferences (2020): https://www.r-project.org/coc.html<br>Code of conduct for UNFCCC conferences, meetings and events (2020): https://unfccc.int/about-us/code-of-conduct-for-unfccc-conferences-meetings-and-events<br>Code of Conduct. Evolution 2020 society for the study of evolution annual meeting June 19-23 Cleveland, Ohio: https://www.evolutionmeetings.org/safe-evolution.html |
| **Example 18** | Society of Ethnobiology Code of Ethics (2019): https://ethnobiology.org/about-society-ethnobiology/ethics<br><br>Principles of Archaeological Ethics-Society for American Archaeology (2020): https://www.saa.org/career-practice/ethics-in-professional-archaeology |
| **Example 19** | Best Practices for Lactation Spaces for Event Organizers: https://medium.com/@jackiekazil/best-practices-for-lactations-spaces-for-event-organizers-8b6c77797c45<br>Add a Lactation Room to Your Checklist: https://ejewishphilanthropy.com/planning-a-conference-add-a-lactation-room-to-your-checklist/<br>How to accommodate a breastpumping mom at your event: https://miriamposner.com/blog/how-to-accommodate-a-breastpumping-mom-at-your-event/<br>Got milk? When packing for a conference requires remembering the breast pump. Science: https://www.sciencemag.org/careers/2018/02/got-milk-when-packing-conference-requires-remembering-breast-pump |
| **Example 20** | A petition by conference attendees to Improve childcare support at future IS-MPMI congresses: https://docs.google.com/document/d/1QAdgecBEig6sgl81aXpk22EIVjdNqWIxevi2r3bzgaM/edit<br>Childcare and Family Resources at The Allied Genetics Conference (TAGC) (2020): https://genetics-gsa.org/tagc-2020/childcare-and-family-resources/<br>Childcare & Nursing at Evolution 2020 meeting, Cleveland, Ohio, USA (2020): https://www.evolutionmeetings.org/childcare--nursing.html<br>Accessibility at I Scientist meeting Technische Universität Berlin (2019): https://www.iscientist.berlin/accessibility |

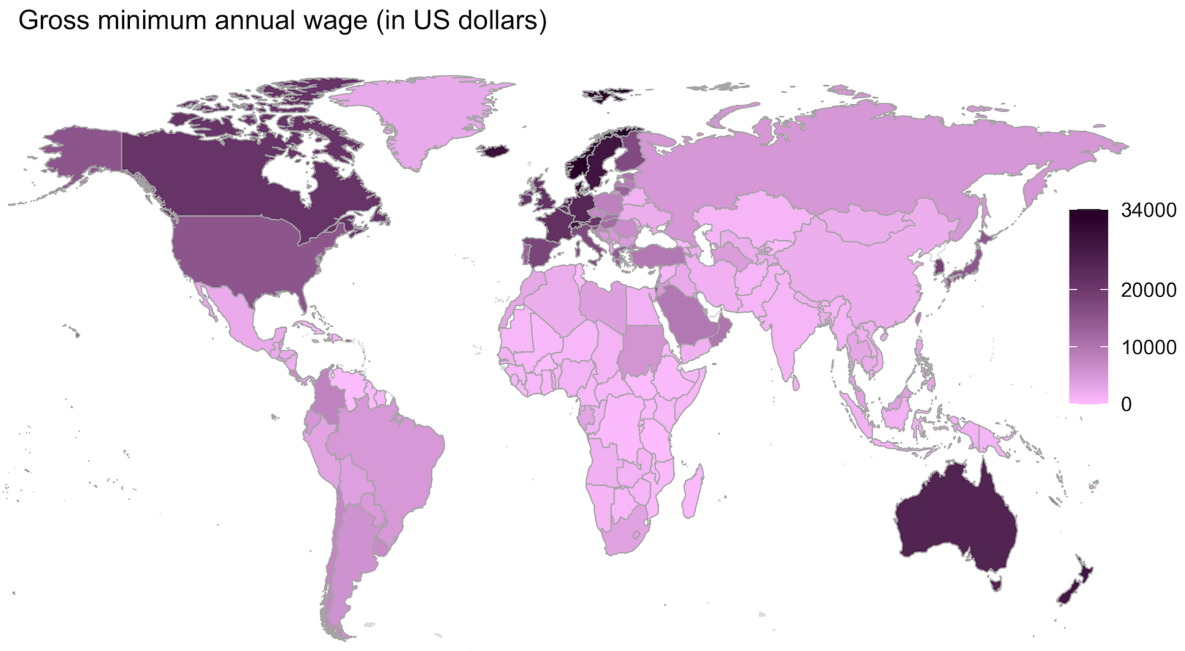


**Figure S1.** Shown is the gross (full amount an employer pays before taxes and other deductions are withheld) minimum annual (12 months) wages (in US dollars) for the year 2019 (data source https://data.worldbank.org/indicator/PA.NUS.PRVT.PP & https://stats.oecd.org/Index.aspx?DataSetCode=RMW). Early career researcher salaries are at minimum annual wage worldwide (https://www.payscale.com/research/UK/Job=Postdoctoral_Research_Associate/Salary). Attending a single national or international conference typically costs USD $1,000-4,000 (https://elifeambassadors.github.io/improving-conferences/) hence attending in-person conferences is not feasible for many researchers in particular early researchers worldwide.

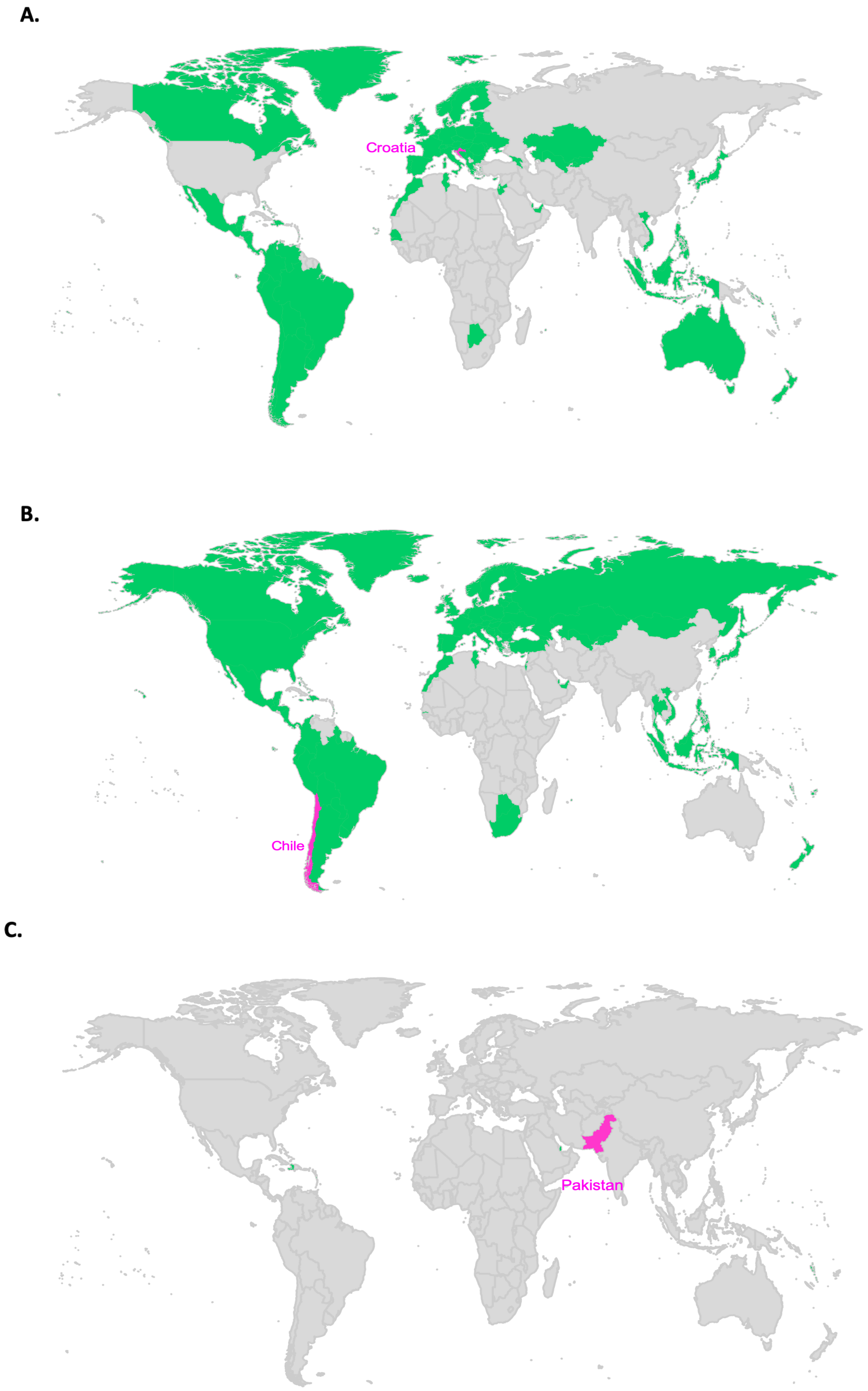
A.
Croatia
B.
Chile
C.
Pakistan

**D.**

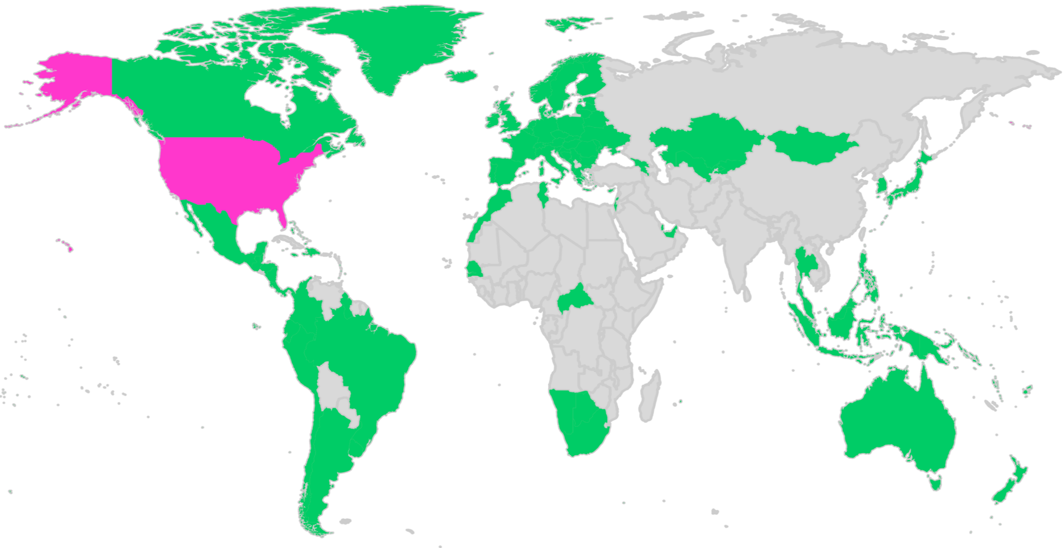

**E.**

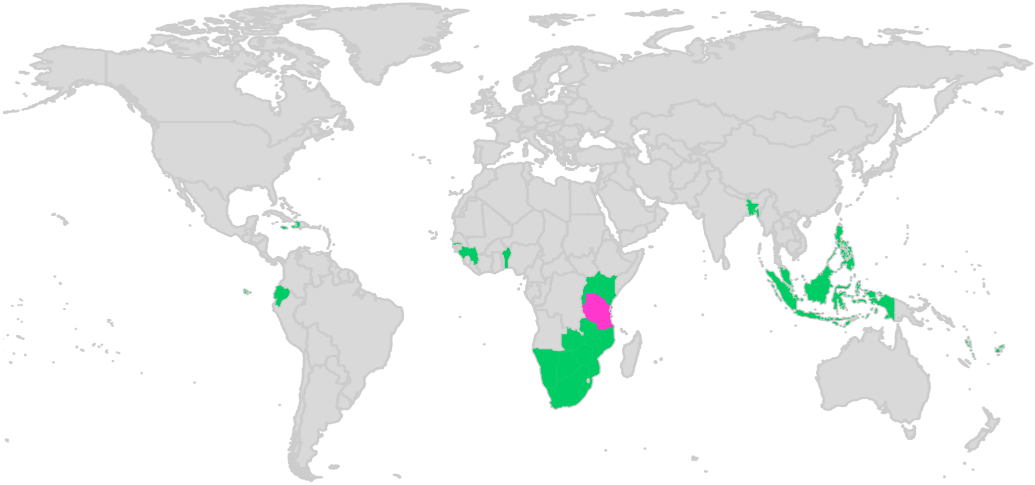

**F.**

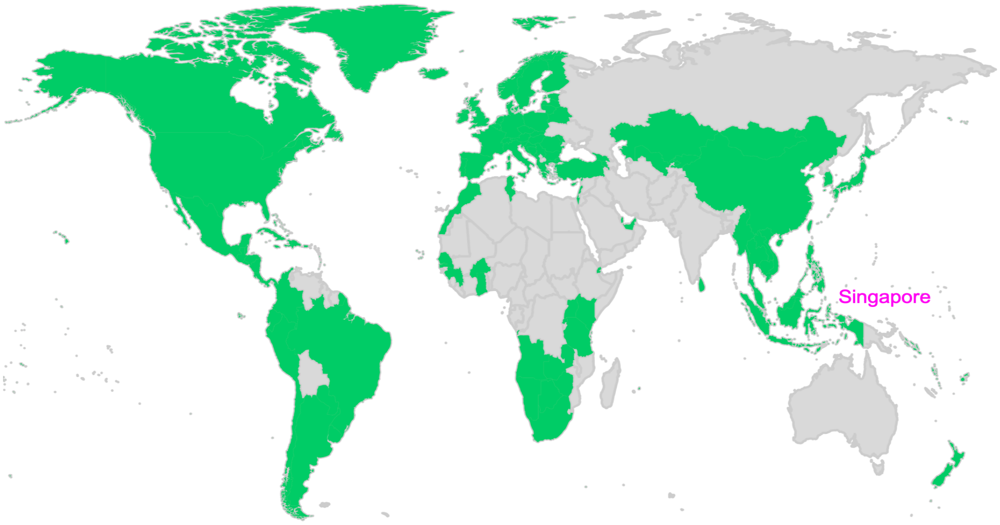

G.

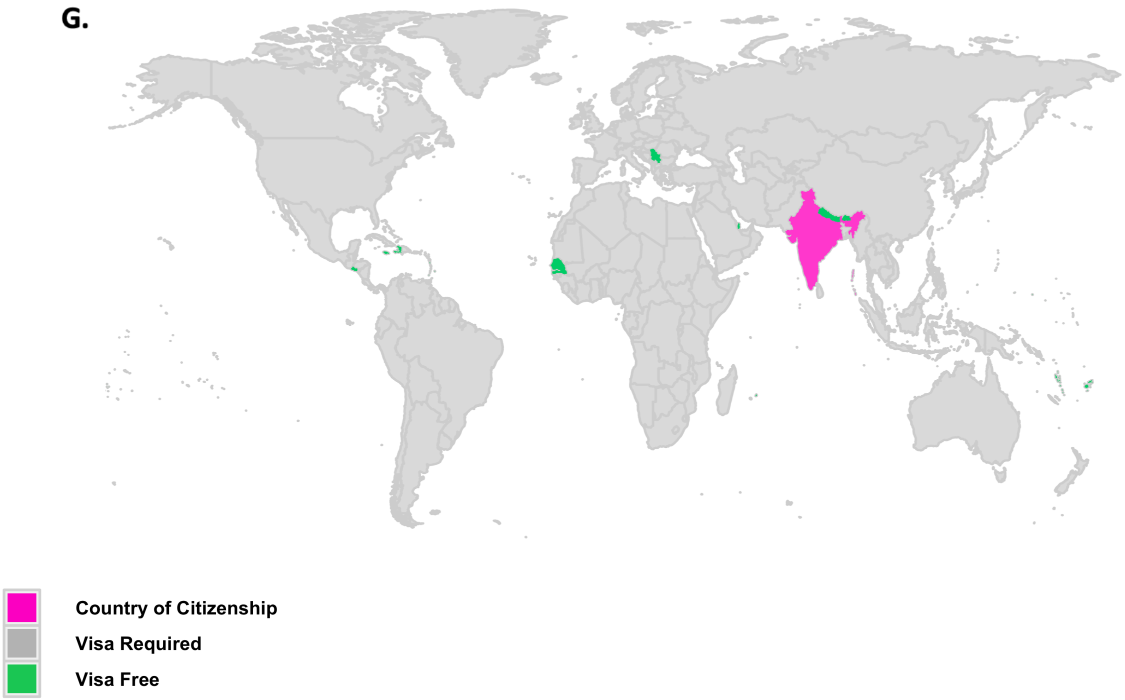


**Figure S2. Visa restrictions and lack of digital conferencing limit scientific career development.** Short-term visitor visa requirements for researchers who are citizens of A) Croatia, B) Chile, C) Pakistan, D) United States E) Tanzania, F) Singapore and G) India. Researchers who are citizens of these countries (in pink) can only travel (short-term for a conference) to select countries (in green) without applying for a visa. Pink color indicates country of citizenship, green color indicates countries researchers can visit without applying for a visa, gray regions indicate that visa required to travel (Data source: The Henley & Partners Passport Index: https://www.henleyglobal.com/henley-passport-index/)

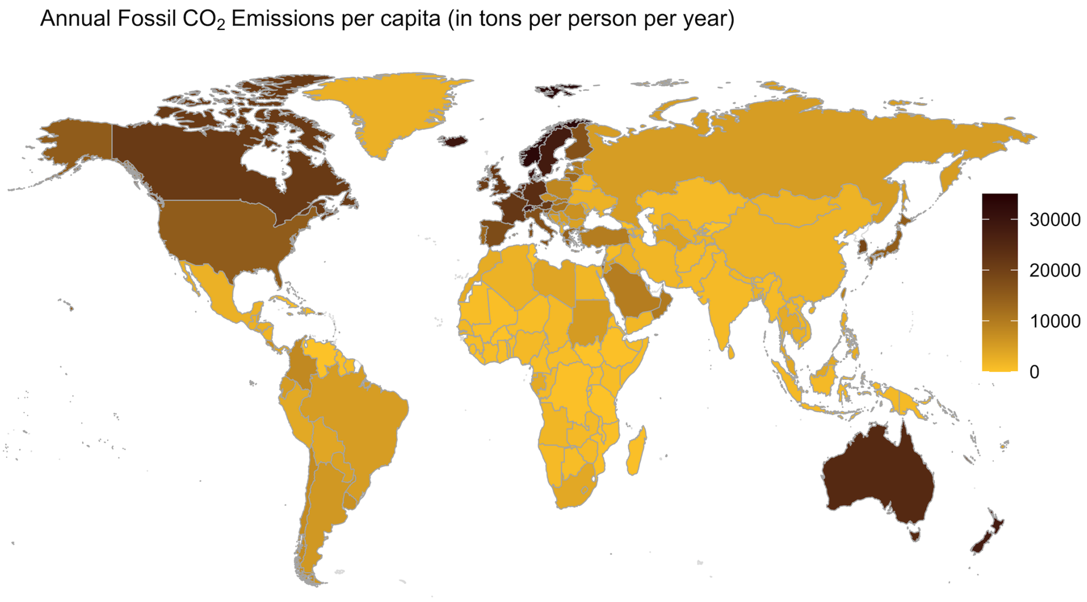


**Figure S3.** The Annual Fossil $CO_2$ Emission per capita: Qatar at 47.8 tons of $CO_2$ per capita, United States at 16.4 tons of $CO_2$ per capita (more than three times the global per capita average), and China at 7.4 tons per capita are amongst the highest emitting countries. Data shown here is for the year 2017 (Data source: https://ourworldindata.org/co2-and-other-greenhouse-gas-emissions & https://www.epa.gov/ghgemissions/global-greenhouse-gas-emissions-data).
